# Standardizing image-derived fish length-frequency distributions to reference measurements using bin-specific error matrices

**DOI:** 10.64898/2026.07.06.736664

**Authors:** Yasutoki Shibata, Yuka Iwahara, Haruhiko Hino, Akiha Tsukada, Yuga Kisara, Tomoya Nishino, Hirotaka Endo

## Abstract

Artificial intelligence (AI)-based image analysis can efficiently estimate fish length, but differences in devices, imaging conditions, operators, and AI models limit comparability among surveys. We propose a standardization framework that estimates a bin-specific error matrix from paired reference measurements and AI-derived lengths and applies it to standardize (correct) AI-derived length-frequency distributions. The Richardson–Lucy expectation-maximization algorithm was used, with the number of iterations selected via cross-validation. Simulations based on empirical length-frequency data from 110 species showed that standardization reduced relative bias and distributional discrepancy; median relative-bias and root mean square error ratios were below 1, and the performance was more affected by the amount of paired data than by the number of cross-validation folds. In real data from 957 Japanese jack mackerel, standardized AI-derived distributions approached human-observer histograms, although discrepancies remained in the range of 160–230 mm. The proposed framework provides a practical approach for improving the comparability of image-derived length-frequency data using paired calibration data, without retraining the underlying AI model.

## Introduction

Recent advances in artificial intelligence (AI)-based image analysis have enabled automated extraction of biological information from fish images across various applications (French et al. 2020; Lu et al. 2020; Lin et al. 2026; Iwahara et al. 2026). In particular, several studies have focused on estimating fish length from images (Hao et al. 2015; Álvarez-Ellacuría et al. 2020; Palmer et al. 2022; Ovalle et al. 2022; Tonachella et al. 2022; Yu et al. 2023; Climent-Perez et al. 2024; Shibata et al. 2024). Several image-based systems have already been implemented in operational or near-operational settings. Palmer et al. (2022) developed and operationalized a deep-learning-based system for estimating fish numbers and mean fork length (FL) from images of landed fish boxes over an entire fishing season. In electronic monitoring, camera-based systems have also been used to collect fish-length data or automate detection, counting, and species classification from fishing videos, although the feasibility and accuracy of automated length estimation can vary among survey designs, imaging settings, and target species.

Fish-length data underpin length-frequency distributions, which are fundamental inputs for stock assessment and growth analysis (Schnute and Fournier 1980; Methot and Wetzel 2013), and improving the efficiency of their acquisition is crucial for research and practical applications (Shibata et al. 2024; Shedrawi et al. 2024). Therefore, advances in image-based length estimation have the potential to substantially enhance data collection systems in fisheries science. In this study, the term AI is used broadly to refer to image-analysis models, including machine– and deep-learning models, which derive fish-length information from images, rather than to generative AI systems.

Various approaches have been proposed for image-based fish length estimation, including methods that use images captured under fixed camera setups (Álvarez-Ellacuría et al. 2020; Palmer et al. 2022; Ovalle et al. 2022; Climent-Perez et al. 2024) and those that rely on mobile devices, such as smartphones and tablets (Shibata et al. 2024; Shedrawi et al. 2024). However, most of these approaches require training AI models using large numbers of annotated images, making the collection and preparation of training data a substantial burden.

This problem becomes particularly critical when integrating data across multiple surveys that differ in devices, AI models, imaging conditions, operators (if mobile devices were used), target species, and fish posture or orientation. In principle, one way to ensure comparability would be to collect sufficient training data for each field condition and develop dedicated imaging and inference systems using well-trained AI models. However, several practical limitations arise when considering deployment across multiple locations and conditions. In some cases, sufficient training data are difficult to obtain owing to differences in regions, fishing methods, or target species. Additionally, budget constraints may require simplification of imaging systems. Consequently, variation in model performance and imaging conditions is inevitable, and the errors associated with AI-derived lengths are unlikely to be consistent across surveys. In this framework, errors in AI-derived lengths are defined relative to the chosen reference measurements, because the true lengths are not directly observed in practical survey settings. Even if it were possible to make model performance and imaging conditions consistent, achieving this would incur substantial development and maintenance costs, potentially resulting in systems that are not sustainable in practice.

Moreover, long-term monitoring and the integration of multiple surveys require that length-frequency data collected under different observational conditions be standardized to a common reference (human measurement) to ensure comparability. Electronic monitoring has already been tested and implemented across a wide range of fisheries and operational conditions (van Helmond et al. 2020). This diversity of observation settings highlights the need for standardization methods that can make length-frequency data comparable across surveys. Herein, we use the term “standardization” in a practical sense to refer to the process of correcting AI-derived length-frequency distributions to make them consistent with a chosen reference.

Two broad approaches can be considered for standardizing AI-derived lengths. The first approach incorporates estimation errors into the training process by retraining the AI model so that it directly outputs standardized lengths. Earlier image-based length-estimation studies have trained models to minimize differences between model outputs and human-provided length observations during training (Ovalle et al. 2022). However, this approach primarily aims to enhance individual-level predictive performance rather than explicitly modeling error structures for post hoc standardization. While this approach can effectively maximize AI performance under a fixed imaging environment at a single site, it becomes excessively costly as the number of survey sites and operating conditions increases, primarily because it requires the use of identical imaging systems and comparable quantities and quality of annotated data.

The second approach compares AI-derived lengths with reference measurements and corrects the AI-derived lengths externally based on their differences, without modifying the AI model itself. Although this approach requires calibration data (described below) to quantify the differences, it does not require identical imaging systems or annotation procedures and can standardize data collected under heterogeneous conditions to a common reference. Therefore, a framework that explicitly models the structure of estimation errors and standardizes length-frequency distributions accordingly is highly practical when the goal is to integrate data from multiple sites, diverse conditions, and long-term surveys.

In this study, rather than directly standardizing continuous length measurements, we discretize length measurements into predefined bins and estimate an error matrix (confusion matrix; e.g., French et al., 2020 for species classification) representing misclassification probabilities between bins based on paired reference measurements and AI-derived lengths. Using this matrix, we propose a framework to standardize (reconstruct) unknown reference-based length-frequency distributions from uncorrected AI-derived length-frequency distributions.

There are two reasons for focusing on length-frequency distributions. First, in fisheries stock assessment, length data are typically treated as categorized data rather than continuous values in the form of length-frequency or length-composition distributions (ICES 2017; Methot et al. 2020; Shibata et al. 2021). Second, estimation errors can be interpreted as probabilistic transitions of individuals across bin boundaries and error matrices provide a direct representation of this structure.

Thus, we focused not on improving individual-level agreement between AI-derived lengths and reference measurements but on developing a methodological framework for standardizing image-derived length-frequency distributions to make them consistent with reference-based (observer-based) length-frequency distributions and comparable across different sites and conditions.

### Objectives

Our objective was to provide a general framework for standardizing AI-derived fish lengths obtained from images into comparable length-frequency distributions by correcting them using a bin-specific error matrix estimated from paired reference measurements and AI-derived lengths. To achieve this objective, the study included four components: (i) development of a method for standardizing AI-derived length-frequency distributions into reference-based length-frequency distributions using an error matrix derived from paired measurements; (ii) simulation-based evaluation of the effects of error magnitude, sample size, and the number of cross-validation folds on standardization performance; (iii) application of the framework to real data to assess the similarity of standardized AI-derived length-frequency distributions to those obtained by human measurers; and (iv) provision of practical guidelines for calibration design and standardization to make image-derived length data comparable across surveys.

## Materials and Methods

The overall workflow of the analysis is illustrated in Fig. 1, and the notation used throughout the study is summarized in Table 1. The Richardson–Lucy algorithm (Richardson 1972; Lucy 1974) was adopted for standardizing length-frequency distributions and applied to the problem setting of this study. The Richardson–Lucy algorithm is an iterative maximum likelihood estimation method originally proposed for image restoration and distribution deconvolution, and it can be interpreted as an expectation–maximization (EM) algorithm under a Poisson observation model (Richardson 1972; Lucy 1974; Shepp and Vardi 1982). Hereafter, we refer to this EM interpretation of the Richardson–Lucy algorithm as Richardson–Lucy EM (RL-EM).

**Fig. 1.**
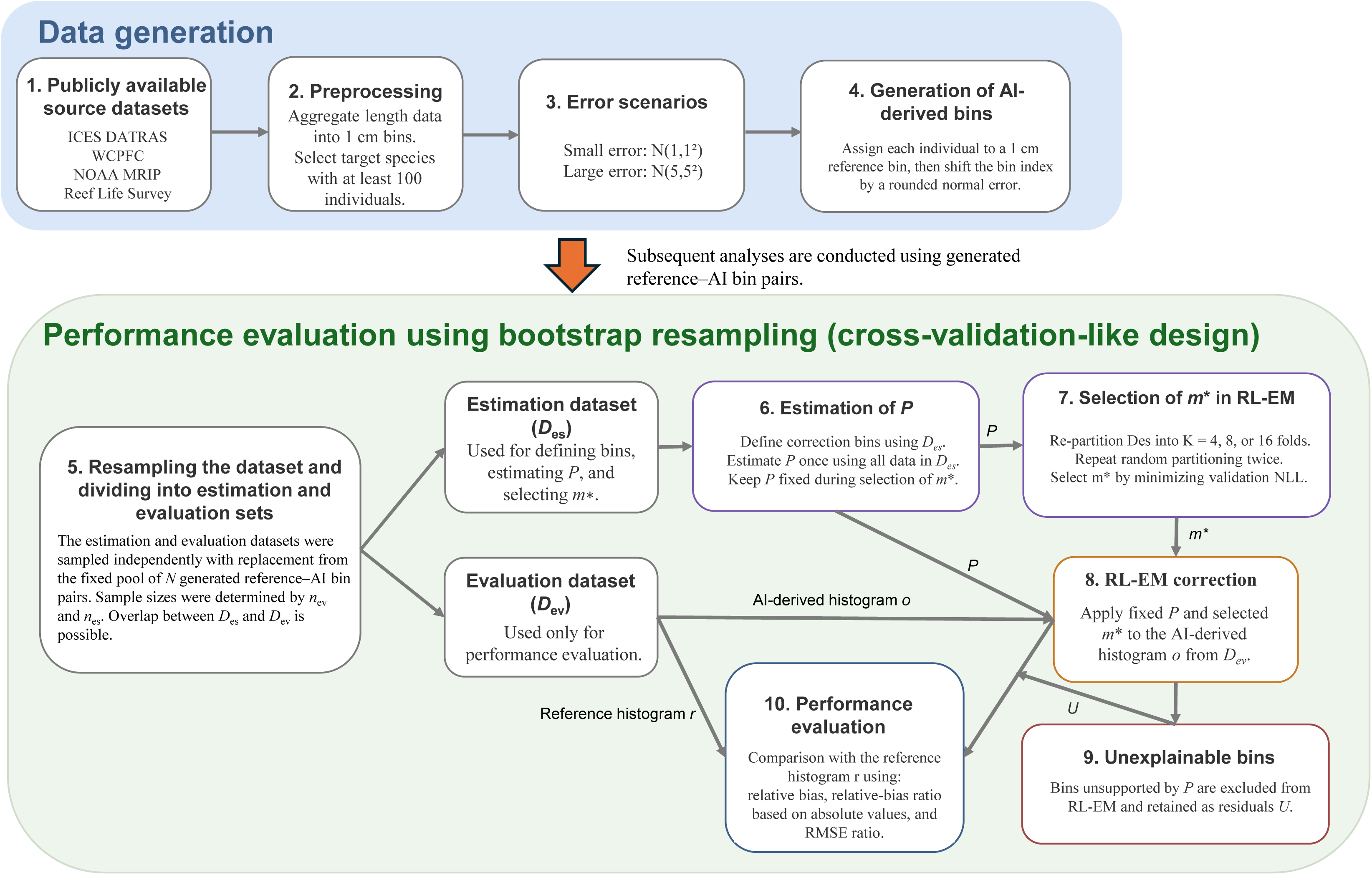
Overview of the simulation workflow, including data generation, error matrix estimation, selection of the number of iterations and evaluation of standardization performance.

**Table 1.**
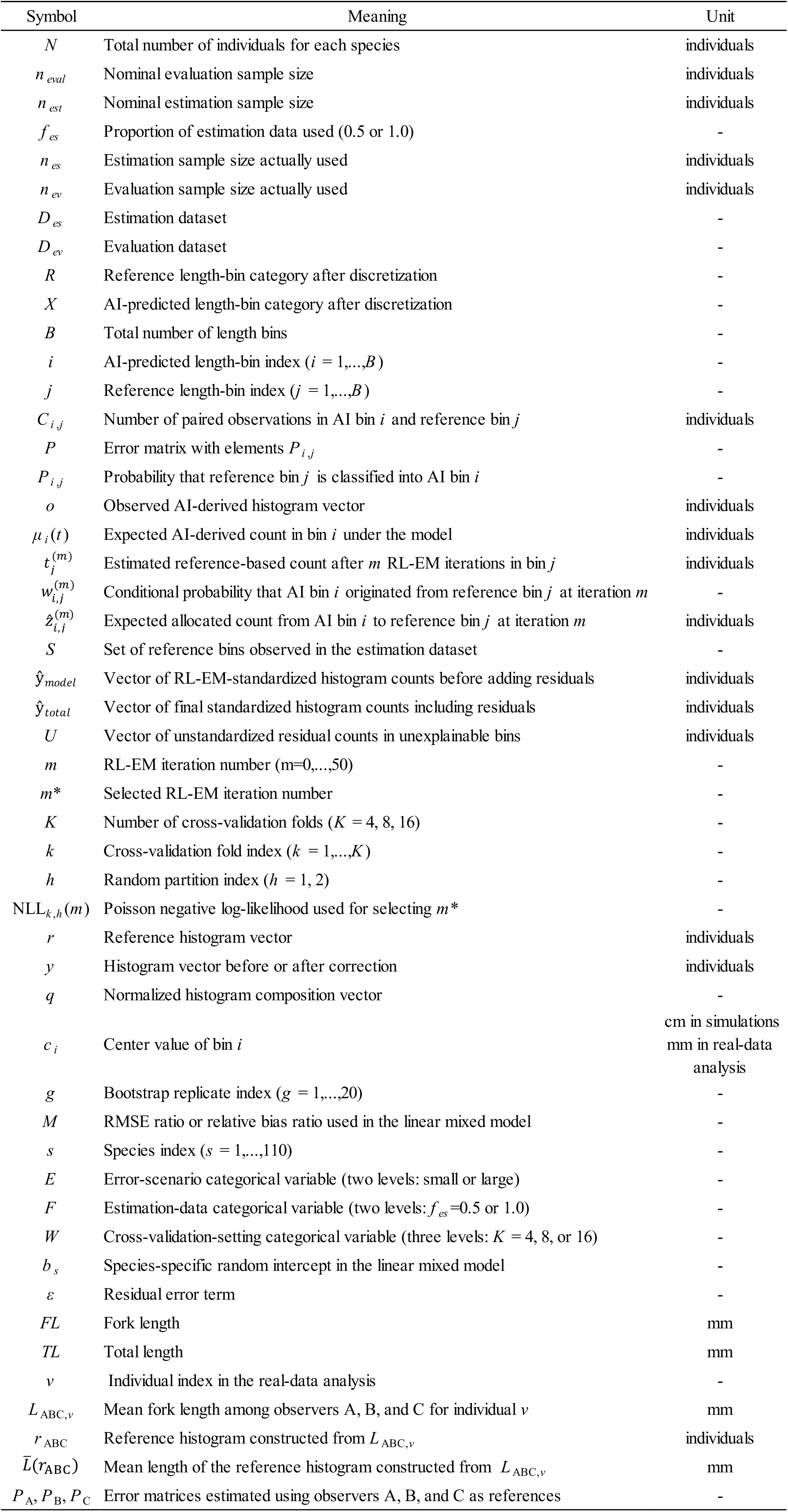
Summary of the notation and symbols used throughout this study, including those related to the RL-EM standardization framework, simulation settings, cross-validation procedures, linear mixed models, and real-data analysis.

In fisheries science, a similar idea of estimating more accurate compositions from surrogate compositions containing errors has already been reported (Hoenig and Heisey 1987). They presented a general framework in which the EM algorithm was used to correct and estimate stock and age compositions from misclassified data, which is conceptually similar to the problem addressed here. In contrast, we assumed an operational framework in which an error matrix *P* was estimated from paired data comprising one-to-one correspondences between reference measurements and AI-derived lengths (Fig. 2), and the number of iterations *m* was selected in a data-driven manner. Following this calibration and tuning step, *P* and the selected iteration number *m*\* were fixed and used to correct the AI-derived length-frequency distribution as described below. This framework enables subsequent measurements to be performed primarily by AI, without requiring reference measurements for every individual. In contrast, Hoenig and Heisey (1987) did not distinguish between the data used for estimation and operation. Instead, fully paired data (with reference values) and incomplete data (without such correspondence) were used jointly to estimate *P* and compositions (Hoenig and Heisey 1987). Therefore, our assumed operational scheme, where *P* is estimated in advance and then fixed for application to new data, has not been considered previously.

**Fig. 2.**
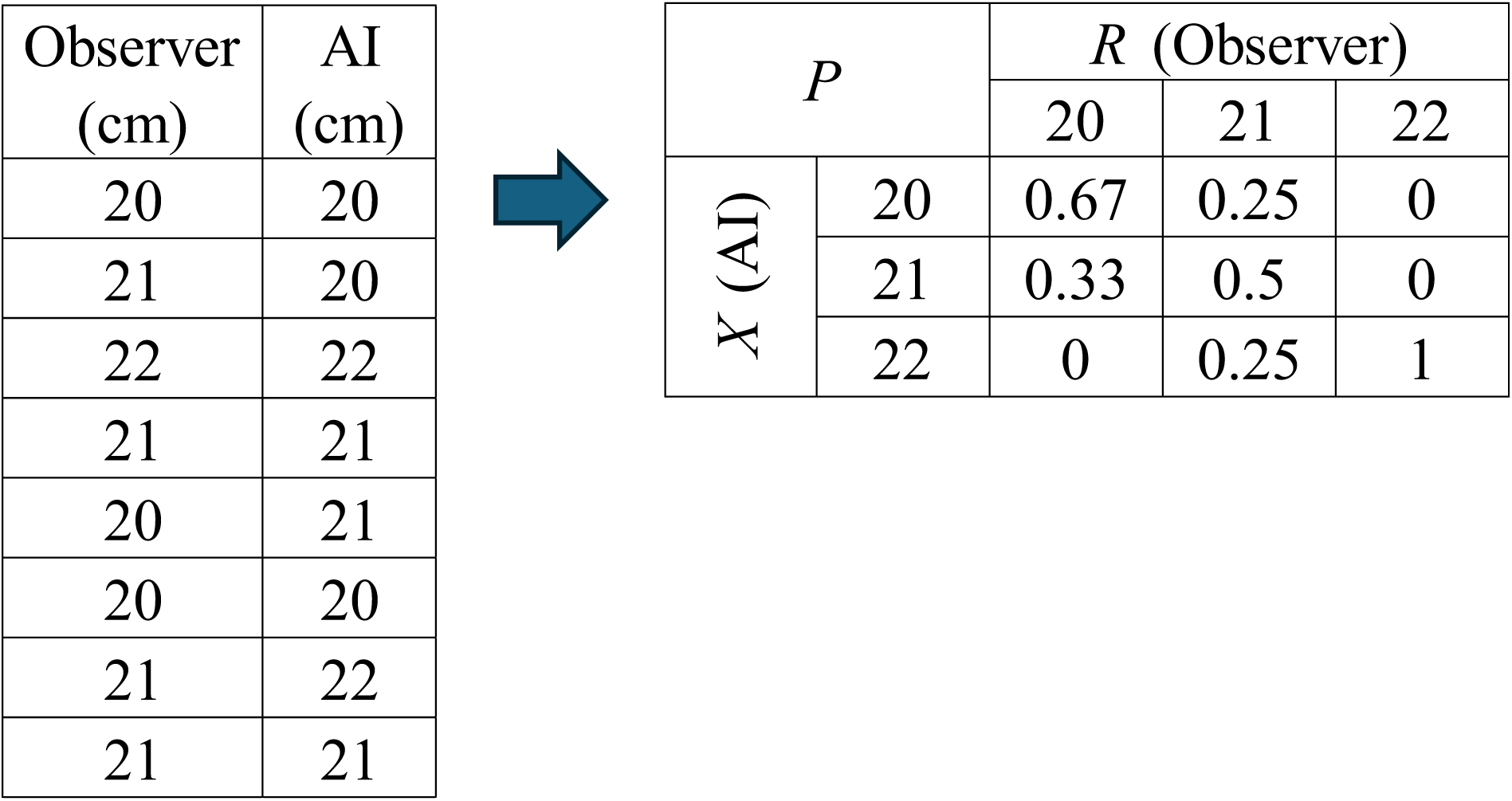
Conceptual diagram for estimating the error matrix *P* from individual pair data. The left panel shows paired data comprising reference measurements and AI-derived lengths for the same individuals. The right panel shows the estimated error matrix *P*, in which the columns represent reference length bins and rows represent AI-predicted length bins. Each column is normalized to sum to one.

### Simulation datasets

To evaluate the performance of our proposed method across multiple real fish species, we used observed length-frequency distributions from existing datasets rather than assuming artificial distributions. The data sources used herein were the International Council for the Exploration of the Sea Database of Trawl Surveys (ICES DATRAS; ICES 2025), the Western and Central Pacific Fisheries Commission (WCPFC 2025), the National Oceanic and Atmospheric Administration (NOAA) Marine Recreational Information Program (MRIP) (NOAA Fisheries 2025), and Reef Life Survey (RLS) (Barrett and Edgar 2015). These datasets were selected to include diverse species, size ranges, and length-frequency distribution shapes. This design was intended to evaluate whether the proposed method performs consistently across empirical length-frequency distributions with different shapes and size ranges.

ICES DATRAS data were obtained from the ICES data portal, comprising data from the North Sea International Bottom Trawl Survey (NS-IBTS) in 2024. These publicly available data are provided under the Creative Commons Attribution 4.0 (CC BY 4.0) license in accordance with the ICES data policy. RLS data were obtained via the Australian Ocean Data Network portal from the dataset “Condition of rocky reef communities around Tasmania: fish surveys” (Barrett and Edgar, 2015; 1992–2007). In this study, only data collected in 2007 were used. These data are also distributed under the Creative Commons Attribution 4.0 (CC BY 4.0) license. NOAA MRIP data in 2024 were obtained using the MRIP query system, which provides open access to recreational fishing data products and query-based estimates. Multiple subsets corresponding to different survey periods (six datasets) were combined and aggregated into species-specific length-frequency distributions. WCPFC data in 2024 were based on publicly available aggregated size data, from which the length-frequency datasets used in this study were constructed. As data acquisition methods and usage conditions differ among databases, the processed length-frequency datasets used in this study, along with the code used to generate them, have been made publicly available in a Zenodo repository (DOI: 10.5281/zenodo.20587024) to ensure reproducibility. The source datasets used for constructing the processed length-frequency datasets were downloaded or queried on November 5, 2025. Length data from all sources were aggregated into 1 cm bins, and species with fewer than 100 individuals were excluded from the analysis. The number of species analyzed from each database was nine for ICES DATRAS, 10 for WCPFC, 58 for NOAA MRIP and 33 for RLS.

AI-derived length-frequency distributions were generated synthetically by shifting the original length bins because the datasets used do not include images. Two error scenarios were considered: a small error scenario, in which bin offsets were drawn from a normal distribution N(1, 1^2^) and a large error scenario, in which bin offsets were drawn from N(5, 5^2^). The magnitude of the bias was therefore defined in units of the bin width (1 cm). Consequently, smaller individuals tended to be subject to larger proportional errors, whereas larger individuals tended to be subject to smaller proportional errors.

Specifically, each individual was first assigned to a 1 cm reference bin. The corresponding AI-predicted bin was then obtained by shifting the reference bin index by an integer offset, which was calculated by rounding a random value drawn from the specified normal distribution. If the shifted bin corresponded to a length below 0 cm, the individual was assigned to the lowest bin, i.e., the 0–1 cm bin. For each species and error scenario, a fixed pool of paired reference and AI-predicted bins was generated once, and all bootstrap replicates were sampled from this same pool. For each bootstrap replicate, the estimation and evaluation datasets were sampled with replacement from this fixed pool of individual pairs. The original observed length-frequency distributions were treated as reference measurements obtained by human observers.

### Design of the performance evaluation

The standardization performance of the proposed method was evaluated for each target species and each error scenario using bootstrap experiments that emulated a cross-validation framework. We define an “individual pair” as a unit comprising one AI-derived length and one corresponding reference measurement for the same individual. Let *N* denote the total number of individuals for each species. The number of evaluation samples, *n*_eval_, is defined as,

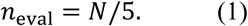

The number of samples used for parameter estimation, *n*_est_, is defined as,

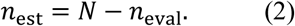

To examine the effect of the sample size used for parameter estimation on standardization performance, the actual number of estimation samples, *n*_es_, is defined as,

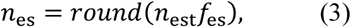

where *f*_es_ represents the proportion of data used for estimation. The value of *f*_es_ was set to 0.5 or 1.0. The number of evaluation samples used in each replicate, *n*_ev_, is defined as,

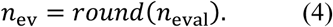

For each *f*_es_ value, 20 bootstrap replicates were performed. Given the large number of species and error scenarios considered, this number of replicates was sufficient to characterize variability in the evaluation metrics while maintaining computational feasibility. In each replicate, the estimation dataset *D*_es_ and the evaluation dataset *D*_ev_ were independently generated by sampling with replacement from the set of individual pairs, with sample sizes *n*_es_ and *n*_ev_, respectively. As sampling was performed with replacement, the same individual pair may appear in both *D*_es_ and *D*_ev_. This reflects the assumption that a common error structure is shared between the estimation and evaluation stages. In other words, the present design assumes that the error structure between AI-derived lengths and reference measurements remains unchanged before and after AI replaces human observers in the measurement process. Notably, this bootstrap design evaluates correction performance under the condition that the estimation and evaluation datasets share the same error structure and does not directly assess extrapolation performance under scenarios in which the error structure itself changes (i.e., domain shift).

### Definition of bins (conversion from continuous to discrete values)

The bins used for correction were defined with a width of 1 cm and were determined independently for each replicate using only the estimation data. Specifically, the upper bound of the bins was set to cover the ranges of the continuous reference measurements and AI-derived lengths in the estimation data. The lower bound was set at 0 cm and the upper bound was extended by adding a sufficient number of bins based on the magnitude of the imposed error (bias and variance), with an additional safety margin to avoid truncation at the upper boundary. Once the bin edges were defined, the continuous reference measurements and AI-derived lengths were discretized into the reference length-bin category (*R*) and the AI-predicted length-bin category (*X*), respectively. Information from the evaluation data was not used to define the bins for parameter estimation.

### Estimation of the error matrix (confusion matrix)

The error matrix *P* is defined such that columns correspond to reference length bins and rows correspond to AI-predicted length bins, as follows:

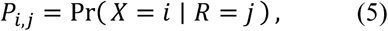

for *i* = 1,…,*B* and *j* = 1,…,*B*, where *B* denotes the number of bins. For each estimation dataset, let *C_i,j_* denote the number of individuals whose reference length falls in bin *j* and whose AI-predicted length is classified into bin *i*. *P* was estimated by normalizing these counts along each column as follows:

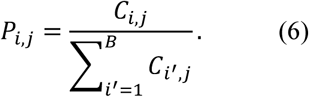

Here, *i*′ is a dummy summation index over AI-predicted bins and is used to avoid ambiguity with the focal index *i* in *P_i,j_*. Thus, each column of *P* sums to one (Fig. 2). No smoothing was applied in the matrix estimation. The set of reference length bins to be corrected was restricted to those that actually appeared in the estimation data. In other words, bins with no observed reference values in the estimation dataset were excluded from the correction.

### Standardization of length-frequency distributions using the Richardson–Lucy EM algorithm

Let *t* = (*t*_1_,…,*t_B_*)^T^ denote the unknown reference-based length-frequency distribution to be standardized (reconstructed). The expected value of the AI-predicted histogram under the error model is given by:

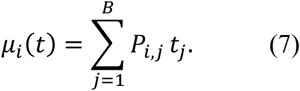

In this study, let *o* = (*o*_1_,…,*o_B_*)^T^ denote the histogram obtained by discretizing AI-derived lengths. The unknown vector *t* is the target of estimation. Under the error model, the observed AI-predicted counts *o_i_* are assumed to follow independent Poisson distributions with means *μ_i_*(*t*). Therefore, *t* can be estimated by maximizing the following log-likelihood:

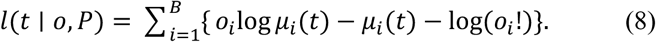

This likelihood provides the basis for the Richardson–Lucy EM update. To derive the update, let the latent variable *z_i_*,*_j_* (described below) denote the unobserved number of individuals in AI-predicted bin *i* that originated from reference bin *j*. The EM algorithm then estimates the expected allocation of observed AI-predicted counts to reference bins and updates *t* accordingly. The term log(*o_i_*!) is a constant independent of *t* and therefore omitting it does not change the maximizer of the likelihood. The EM algorithm can then be expressed as described in two steps.

### E-step

Given the estimate *t*^(*m*)^ at iteration *m*, the conditional probability that an individual observed in AI-predicted bin *i* originates from reference bin *j* is defined as,

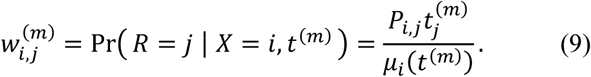

The expected number of individuals allocated from AI-predicted bin *i* to reference bin *j* is then given as,

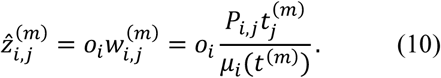

### M-step

The length-frequency distribution was updated by summing the expected counts assigned to each reference bin *j* over all AI bins:

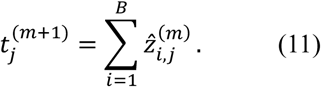

Combining the E-step and M-step gives the following update rule:

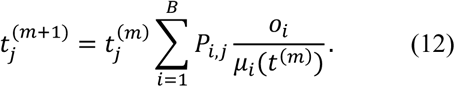

The initial value was set to be uniform over the target bins, with the total sum equal to that of the observed histogram.

### Treatment of unexplainable bins in the standardization procedure

Only length bins in which reference observations were present in the estimation data were included as targets for standardization. Therefore, among the AI-predicted bins *i* in the evaluation data, those for which the total assignment probability from all target reference bins was zero in *P* estimated from the estimation data were defined as unexplainable bins. That is, among AI-predicted bins *i* in the data to be corrected, bins satisfying

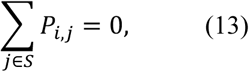

were classified as unexplainable, where *S* denotes the set of reference bins observed in the estimation data. This situation arises when AI-predicted values fall into length ranges that were not observed in the estimation data. Individuals falling into unexplainable bins were not used to terminate the estimation procedure. Instead, they were excluded from the RL-EM input and retained as unstandardized residuals. Accordingly, the output length-frequency distribution after applying the standardization procedure is defined as

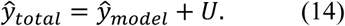

Here, *y*^_model_ denotes the RL-EM estimate *t*^(^*^m^*^*)^ after the selected number of iterations, as described below, and represents the component standardized by RL-EM. The vector *U* represents the unstandardized residual component retained outside the RL-EM procedure. This formulation does not imply that the residuals were reintroduced into the RL-EM procedure. Rather, it indicates that individuals classified as unexplainable were added to the standardized component after RL-EM estimation. A simple example of the standardization process using *P* is provided in Supplementary 1.

### Selection of the number of iterations *m*

The RL-EM algorithm was not iterated indefinitely; instead, the number of iterations *m* was selected in a data-driven manner. Richardson–Lucy deconvolution is known to amplify noise and become unstable as *m* increases, and in practical applications, early stopping of iterations functions as a form of regularization (Prato et al. 2012). In the present setting, *P* itself is estimated from a finite set of paired data. Therefore, excessive iterations may cause estimation error in *P* and sampling variability in the data to be overly reflected in the AI-predicted histogram. Therefore, *m* was selected using cross-validation within the estimation data. This selection of *m* is not merely a computational convenience; it serves as an effective regularization mechanism to prevent overfitting to the estimation data and to obtain more stable standardized results (Stanley 1995).

The candidate values of *m* were set to *m* = 0,…,50. The number of cross-validation folds *K* was set to 4, 8, or 16 and its effect was evaluated. For each *K*, two independent random re-partitions were performed. For each validation fold, RL-EM was applied for each candidate *m*. For example, when *K* = 4, each candidate *m* was evaluated over 4 x 2 = 8 validation folds. The Poisson negative log-likelihood obtained for each fold was averaged for each candidate *m*, and the value *m\** that minimized the average negative log-likelihood was selected. That is, *m\** is determined as shown,

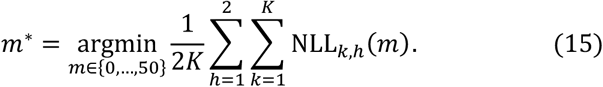

Here, NLL*_k_*,*_h_*(*m*) denotes the Poisson negative log-likelihood for candidate *m* obtained in the *k*-th validation fold of the *h*-th random partition. *P* was estimated once using the full estimation dataset before cross-validation and was kept fixed throughout the selection of *m*, such that only the iteration number was optimized, while the error structure remained unchanged. This design isolates the selection of *m* from the estimation of *P* and avoids additional instability due to repeated estimation of *P* within folds, particularly for species with limited sample sizes. Thus, this procedure should be interpreted as cross-validation-based tuning of the iteration number conditional on a fixed *P*, rather than as full cross-validation of *P* and *m*.

For each candidate value of *m*, the same RL-EM update described above was applied to obtain *t*^(*m*)^ within each validation fold. The likelihood in Eq. (8) defines the Poisson observation model underlying this update and was used to derive the RL-EM algorithm. In contrast, the Poisson negative log-likelihood below was used only as a validation criterion for selecting the stopping iteration *m*\*, not as a separate EM update.

For each *m*, the Poisson negative log-likelihood between the RL-EM estimate *t*^(m)^ and the corresponding reference histogram *r* = (*r*_1_,…,*r*_B_)^T^ was calculated as shown,

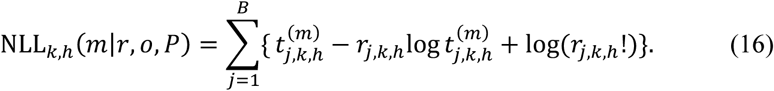

The *m*\* value was then selected as the value that minimized the average negative log-likelihood across all folds and repetitions. The term log(*r_j,k,h_*!) is independent of *m* and therefore omitting this term does not affect the selection of *m*\*. When unexplainable bins occurred within validation folds, the Poisson negative log-likelihood for selecting *m* was calculated using only the explainable component used as input to RL-EM.

### Performance metrics

Performance before and after standardization was evaluated separately for both the estimation and evaluation datasets. Let *r* = (*r*_1_,…,*r*_B_)^T^ denote the reference histogram and *y* = (*y*_1_,…,*y_B_*)^T^ denote the histogram to be compared. The corresponding composition probability vector *q* is defined by normalizing each histogram by its total count as follows:

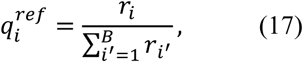

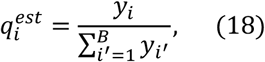

where *y* corresponds to the histogram being evaluated (i.e., *y* = *o* before correction and *y* = *y*^_total_ after correction) and *i* = 1,…, *B*. The root mean square error (RMSE) between the composition probability vectors is defined as,

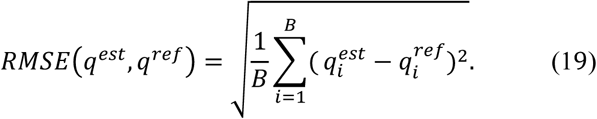

The mean length is defined using the bin centers *c_i_* as follows:

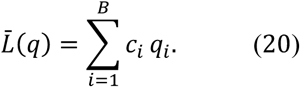

Bias is defined as

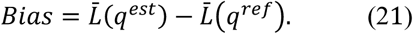

Relative bias (RB) is defined as

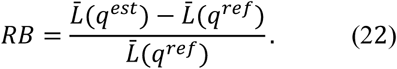

RB was evaluated directly because it reflects both the magnitude and direction of deviation (i.e., overestimation or underestimation). To assess improvement in the magnitude of RB, the RB ratio was calculated as the absolute relative bias after correction divided by the absolute RB before correction. In contrast, the RMSE is a non-negative measure without directional information, and therefore the RMSE ratio was calculated as the RMSE after correction divided by the RMSE before correction. The results of each replicate were aggregated by species, error scenario, proportion of estimation data and number of cross-validation folds and improvements before and after standardization were compared.

### Analysis of factors affecting correction performance

Linear mixed models were fitted using the evaluation metrics obtained from each bootstrap replicate to evaluate which simulation settings affected correction performance. The response variable *M* was either the RMSE ratio or the RB ratio, both calculated from the evaluation dataset. Error-scenario (*E*), estimation-data (*F*) and cross-validation-setting (*W*) categories were treated as fixed effects, and species *s* was treated as a random effect through the species-specific random intercept *b_s_*. The analysis was performed to assess the relative influence of simulation settings on correction performance.

## Real-data analysis

### Positioning of the real-data analysis

In the simulation analysis, the performance characteristics of the proposed method and the effects of standardization conditions were evaluated using evaluation data that were not used for parameter estimation. By contrast, the real-data analysis aimed to demonstrate the extent to which AI-derived length-frequency distributions can be standardized under realistic observational conditions, using paired data obtained from actual measurements to estimate *P*. This analysis is intended to illustrate the behavior of the standardization results under operational conditions after calibration, not to evaluate performance on data not used for calibration, because evaluation under the simulation design was assessed separately. This analysis represents an idealized calibration scenario rather than an independent evaluation of performance on new data.

A separate evaluation dataset was not reserved for the real-data analysis because the number of available paired observations was limited to 957 individuals, and holding out part of the data would have reduced length-bin coverage for estimating the error matrices. Therefore, this analysis was designed to illustrate the standardization behavior when all available paired observations were used for calibration, rather than to provide an independent assessment of generalization performance. Accordingly, for each reference measurer, *P* was estimated using all available paired data from all individuals, and the standardization was applied to the AI-derived length-frequency distribution obtained from the same full dataset. The iteration number *m*\* was also selected through cross-validation within the same dataset. This design reflects a practical scenario in which all available calibration data are used to estimate *P*. Therefore, the results should be interpreted as the level of consistency achievable under standardization based on *P* estimated from the same dataset, rather than as performance expected for independent future datasets.

## Preparation of the real dataset

### Japanese jack mackerel used for measurement error evaluation

A total of 959 Japanese jack mackerel (*Trachurus japonicus*) landed in 2025 in Kagoshima, Shizuoka, and Kanagawa Prefectures in Japan were used to evaluate the differences between measurements obtained by human observers and AI. Japanese jack mackerel is an important target species in Japanese coastal fisheries, and length-frequency information is routinely collected for this species in stock assessment (Yasuda et al. 2025). Following the preprocessing described below, 957 individuals were retained for the real-data analysis. The retained individuals covered a fork-length range of 46–274 mm after preprocessing. All individuals were fresh fish with no history of freezing and were purchased within 1–2 days after landing. They were stored under refrigerated conditions until measurement, and all procedures from imaging to manual measurement by human observers were completed within half a day.

### Imaging and AI-based measurement

A modified version of the smartphone application ToroCam (Shibata et al. 2024) was used for image acquisition. An external camera (EPOS S6 4K Model DSWD1, EPOS GROUP A/S, Denmark) was connected to a smartphone on which ToroCam was installed, and images of fish were captured from above using a camera system fixed on a table (Figs. 3a and 3b). The modified version of ToroCam used in this study is not currently publicly available; however, the imaging setup and data acquisition procedure are described in detail to allow replication. The analysis code is publicly available in a Zenodo repository. During imaging, the device was not touched directly, and a remote shutter was used to avoid camera shake. Fish were placed in a container to prevent individuals from overlapping, and each individual was assigned an identification ID, which was recorded simultaneously in the image (Fig. 3c). All individuals were positioned with their heads oriented to the left, and the caudal fins were arranged as consistently as possible. A keypoint estimation model (described below) was then applied to estimate the FL of each individual.

**Fig. 3.**
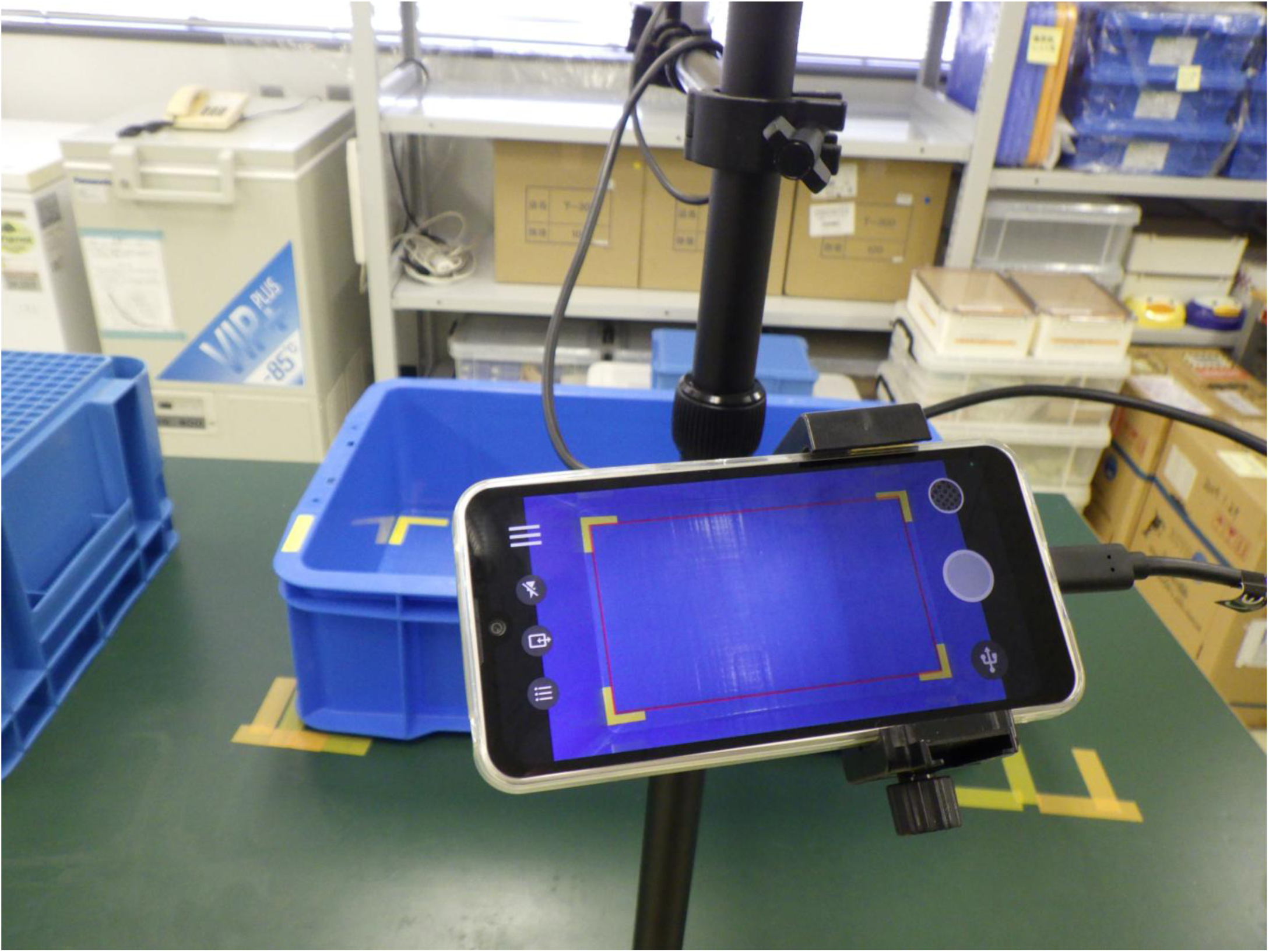
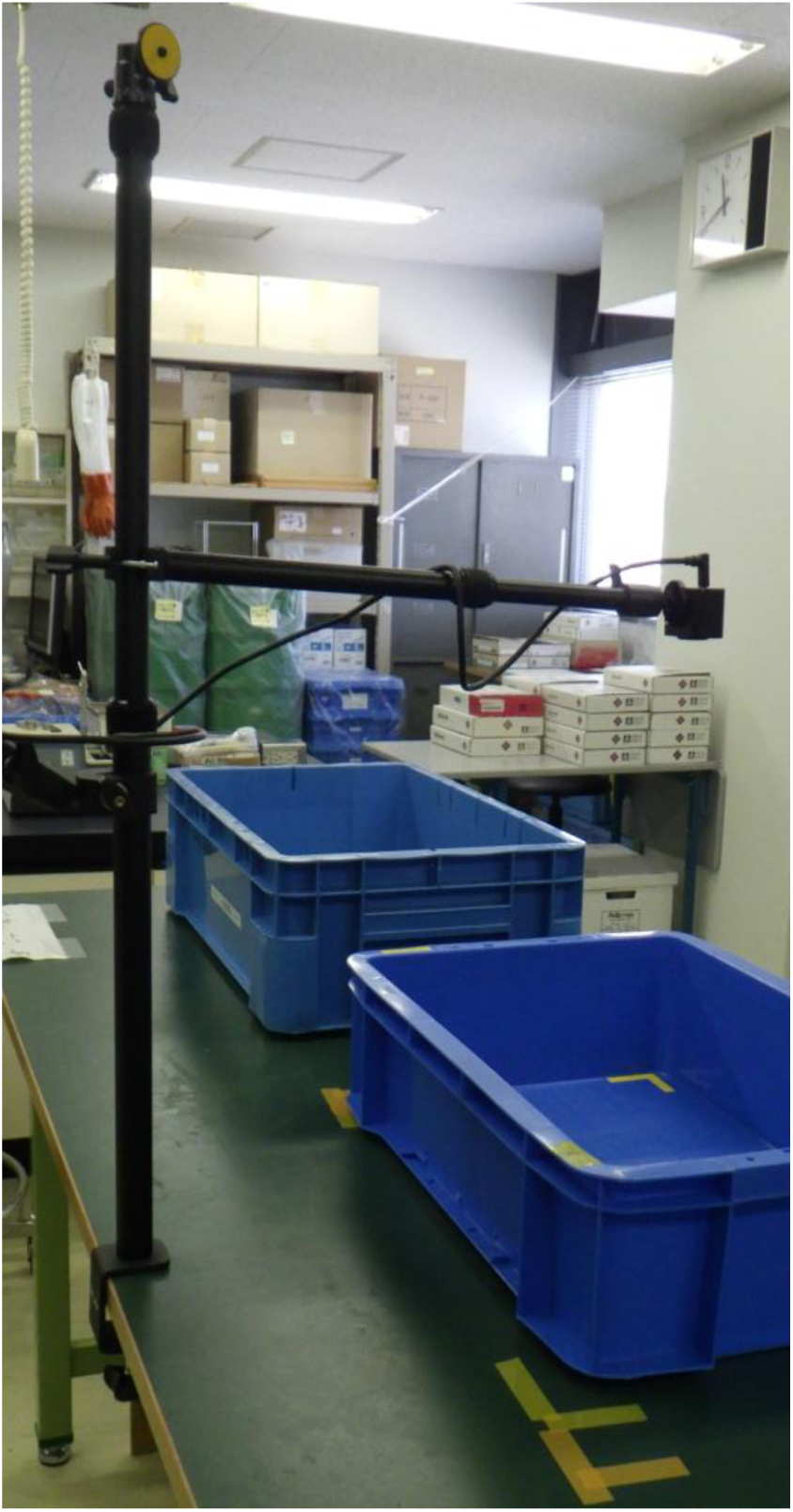
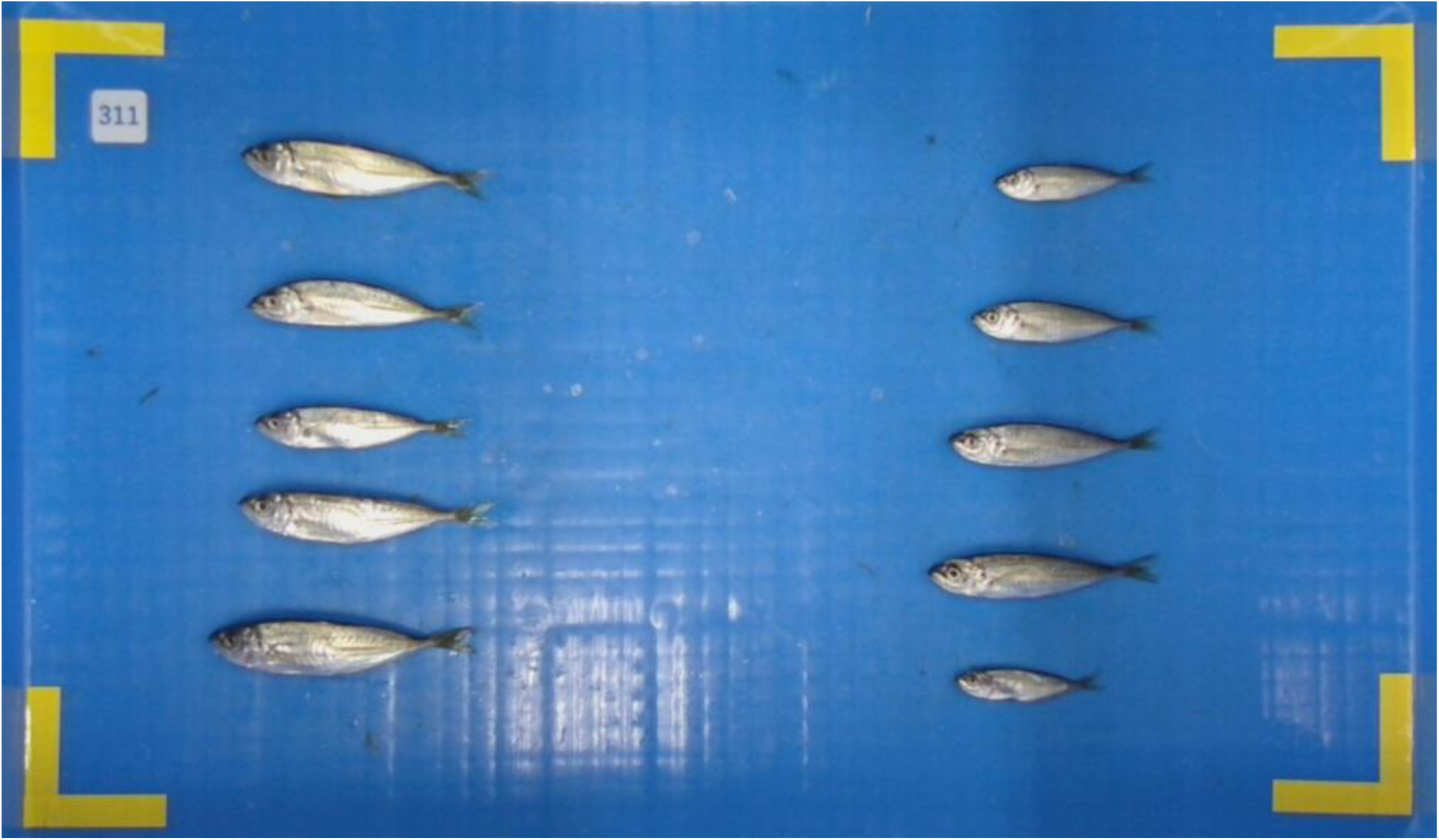
Imaging system and examples used for real-data acquisition. (a) Configuration of the imaging system using ToroCam. (b) Example of fish arrangement during imaging. (c) Example of an image with individual identification IDs.

### Manual measurements by human observers

Following imaging with ToroCam and the external camera, the FL of each individual was measured once independently by three experienced human observers using measuring boards with 1 mm graduations, without knowledge of each other’s measurements. Thus, three independent observations (references) were obtained for each individual. This design enables inter-observer variability to be quantified and provides a reference level for evaluating the consistency of the standardized AI-derived measurements.

### Training data and keypoint estimation model

To obtain AI-derived FLs, a keypoint estimation model was trained on fish images collected at multiple locations between 2020 and 2024 (*n* = 27,016 individuals). Only the total number of individuals in the training dataset is reported because the model was trained only for length estimation and did not use species labels. To stably capture body shape information required for fork length estimation, nine keypoints were defined along the contour of the fish body from the mouth to the caudal fin (Fig. 4). These keypoints were used to constrain the overall body shape and fork length was calculated as the Euclidean distance between keypoints 1 and 7. The dataset was split into training, validation, and test sets at a ratio of 8:1:1, a practical split used in supervised deep-learning workflows and also adopted in previous fish-image length-estimation work (Shibata et al. 2024). The model was trained using HRFormer (Yuan et al. 2021) implemented in the Python package MMPose (MMPose Contributors 2020). Note that the primary objective of this study is not the development of a high-performance AI model itself, but the development of a framework for correcting AI-derived lengths. Therefore, detailed comparisons of different model architectures or training conditions were not conducted. The trained model was applied to images of Japanese jack mackerel collected for the real-data analysis to obtain AI-derived FLs. For keypoint estimation, prior detection of the fish body region is required. This was performed using a previously developed model that detects non-overlapping individual fish (Shibata et al. 2024).

**Fig. 4.**
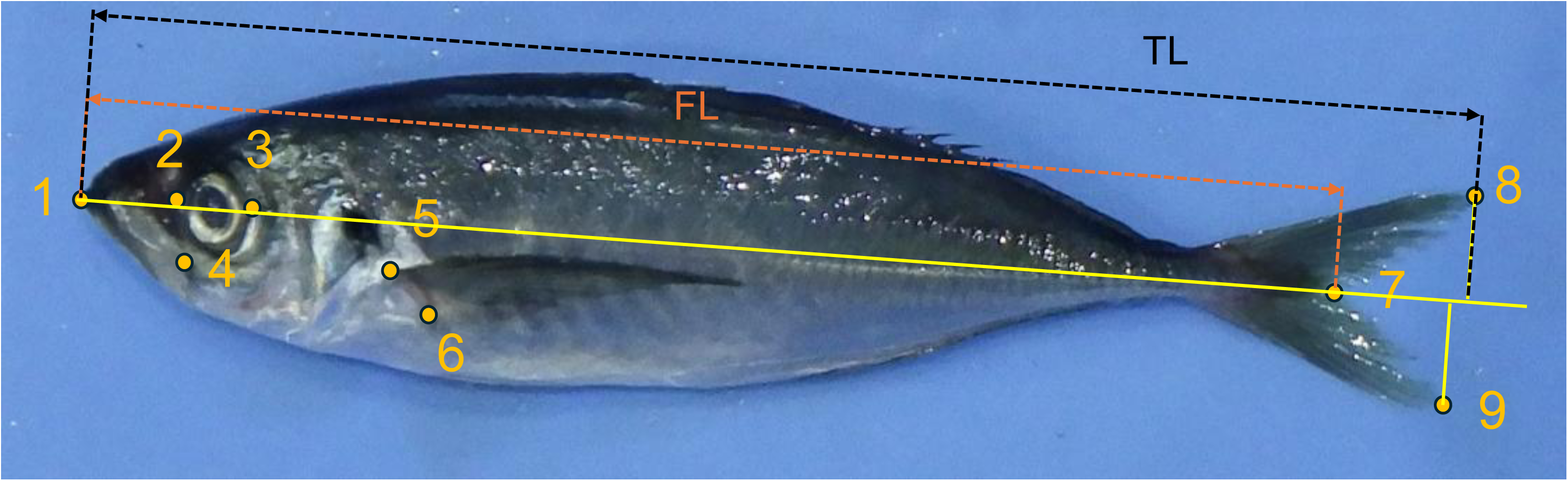
Definition of keypoints for capturing fish body shape. Nine keypoints were defined along the contour from the mouth to the caudal fin. These keypoints were used to constrain the overall body shape. Fork length (FL) was calculated as the Euclidean distance between keypoints 1 and 7. For the FL/TL screening, total length (TL) was approximated as the Euclidean distance from keypoint 1 to the perpendicular projection of keypoint 8 onto the line connecting keypoints 1 and 7.

The 957 Japanese jack mackerel used in the real-data analysis were not included in the training, validation, or test datasets used to develop the keypoint estimation model. The trained model was applied to these images only to obtain AI-derived FLs, which were then treated as fixed inputs for the downstream standardization analysis. Thus, the present study did not retrain, fine-tune, or compare AI models, but instead evaluated a framework for transforming existing AI-derived lengths into length-frequency distributions consistent with reference measurements. The AI-derived lengths, reference measurements, and analysis code are available in the public repository, whereas the image dataset used for model training is not publicly available.

### Real dataset and preprocessing

For the real-data analysis, an integrated dataset was used that included individual IDs, measurer identities, FL measurements by human observers and AI-derived FLs. As a preprocessing step, outliers were removed based on the ratio FL/TL, where TL denotes the total length calculated from the same set of keypoint outputs.

Individuals with FL/TL < 0.8 were excluded. This threshold was determined based on inspection of the empirical distribution to remove clearly implausible values. This ratio was used as a simple geometric consistency check of the keypoint outputs. Additionally, only individuals for whom measurements from all three observers (A, B, and C) and AI-derived lengths were available were included in the analysis. Following preprocessing, the final dataset contained 957 individuals.

### Definition of bins

The bin width used for standardization was set to 5 mm, corresponding to the 5-mm interval used in punching sheets for our routine FL measurements of Japanese jack mackerel. In the real-data analysis, a single common set of bin edges was used for all reference measurers. The bin edges were defined to cover the AI-derived FLs and the measurements from all three human observers. The lower bound was set to include 0 mm and the upper bound was extended by adding five additional bins. All other aspects of bin handling, defining *P*, standardization using RL-EM, the treatment of unexplainable bins and the definition of performance metrics were the same as those used in the simulation analysis.

### Estimation of error matrices for each reference measurer

To compare the degree of standardization achieved under different reference standards, three error matrices, *P*_A_, *P*_B_, and *P*_C_, were constructed using measurers A, B, and C as the reference, respectively. Each error matrix was estimated using all 957 paired observations. Specifically, for each reference measurer, the human-measured FLs were discretized using the common 5-mm bins and treated as the reference length-bin categories *R*. The corresponding AI-derived FLs for the same individuals were discretized using the same bin edges and treated as the AI-predicted length-bin categories *X*. The corresponding error matrix was then estimated from these paired bin categories using the same procedure as in the simulation analysis.

### Selection of the number of iterations *m*

In the real-data analysis, the number of RL-EM iterations *m* was also selected via cross-validation. Candidate values were set to *m* = 0,…,50, and a 4-fold cross-validation was repeated twice. In this case, the error matrices *P*_A_, *P*_B_, and *P*_C_, each estimated using all individuals, were kept fixed, and *m* was selected based on the Poisson negative log-likelihood. This procedure ensures that the selected iteration number corresponds to the final error matrix used in practical applications.

### Calculation of standardized length-frequency distributions

The AI-derived length-frequency distributions were standardized using the error matrices *P*_A_, *P*_B_, and *P*_C_, each estimated with a different reference measurer. As in the simulation analysis, only bins in which reference observations were present were included as targets for correction. Individuals falling into unexplainable bins were not used to terminate the computation, but were excluded from the RL-EM input and retained as unstandardized residuals. Accordingly, the final standardized length-frequency distribution was defined as the sum of the RL-EM-standardized component and the unstandardized residuals.

### Evaluation of standardization performance

The differences among the three human observers were used as a baseline to evaluate the degree of standardization. First, for each *v* (*v* = 1,…,957), the mean of the three measurements is defined as,

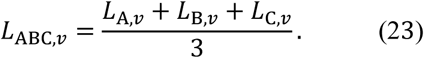

Since the true length-frequency distribution is not available, these mean FLs were used to construct a reference histogram that does not depend on any single measurer. Specifically, *L*_ABC,*v*_ was aggregated into 5-mm bins to obtain the reference histogram *r*_ABC_ = (*r*_ABC,1_,…,*r*_ABC,*B*_)^T^, where *r*_ABC,*i*_ denotes the count in bin *i*. The mean FL of this reference histogram is then defined as,

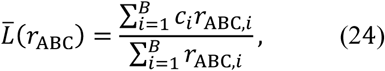

where *c_i_* denotes the center value of bin *i* and *r*_ABC,*i*_ denotes the count in bin *i* of the reference histogram. Histograms were also constructed for the measurements obtained by each observer (A, B, and C), the uncorrected AI-derived histogram, and the AI-derived histograms corrected using each error matrix (*P*_A_, *P*_B_, and *P*_C_). These histograms were compared with *r*_ABC_ to evaluate both inter-observer variability and the degree to which AI-derived histograms approached this common reference after standardization. RB was calculated by comparing the mean FL of each histogram with *Lˉ*(*r*_ABC_), and the RMSE between the corresponding composition probability vectors was calculated.

In the real-data analysis, after/before ratios were not used as the primary metrics. The aim was not to quantify how the AI-derived histogram improved after correction, but to evaluate whether the standardized AI-derived histogram approached the range of variability among human observers. As the Japanese jack mackerel samples were purchased by size category, three distinct peaks were observed in the length-frequency distribution. Therefore, the same metrics were also calculated for three length ranges corresponding to these peaks: 0–120 mm, 160–230 mm and 230–300 mm. For each range-specific analysis, histograms were first restricted to the corresponding length range and then converted to composition probability vectors within that range. RB was calculated from the mean FL derived from these range-specific composition vectors, and the RMSE between the corresponding range-specific composition vectors was calculated.

## Results

### Simulation analysis

In the simulation analysis, correction performance was compared under two error scenarios (*E*; small and large error), two proportions of estimation data (*f*_es_ = 0.5 or 1) and three cross-validation settings (*K* = 4, 8, and 16). Performance was evaluated using the RB after correction, the RB ratio calculated using absolute RB values, and the RMSE ratio, all calculated on the evaluation dataset (*D*_ev_).

The RB after correction converged to approximately zero under all conditions. In the small-error scenario, the median was approximately 0.00%. Even under the large-error scenario, the median remained small at approximately 0.14%, showing that the remaining RB was small in the simulation setting (Fig. 5a).

**Fig. 5.**
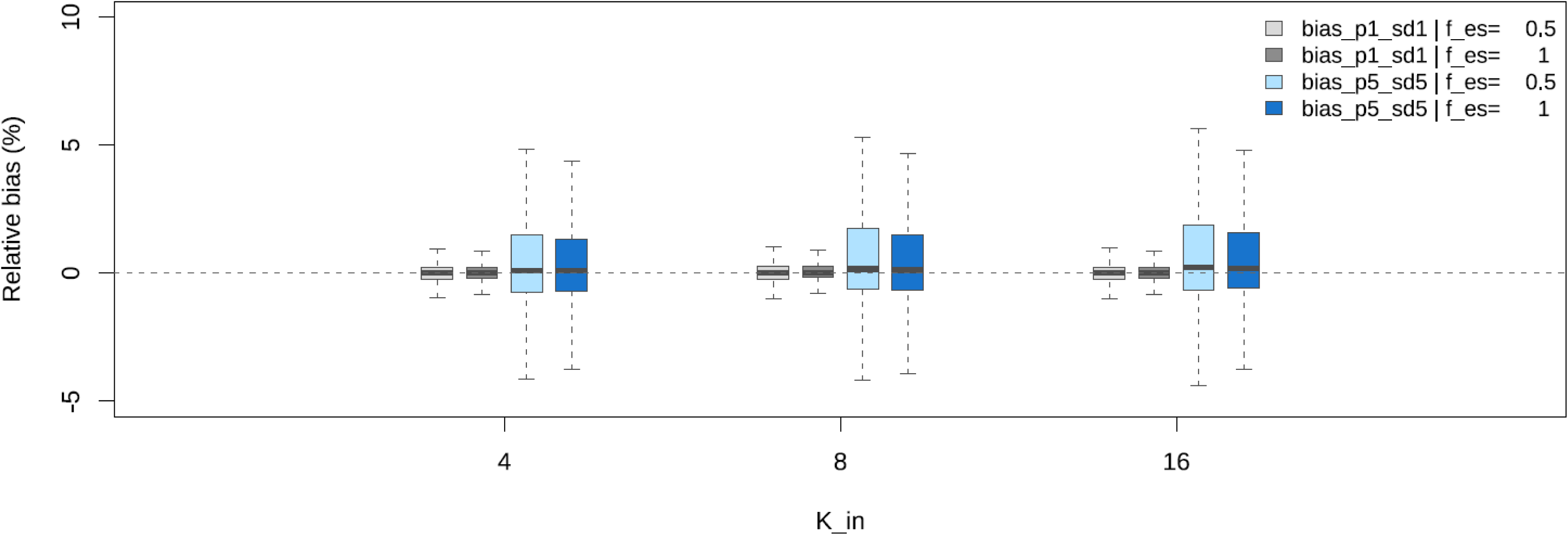
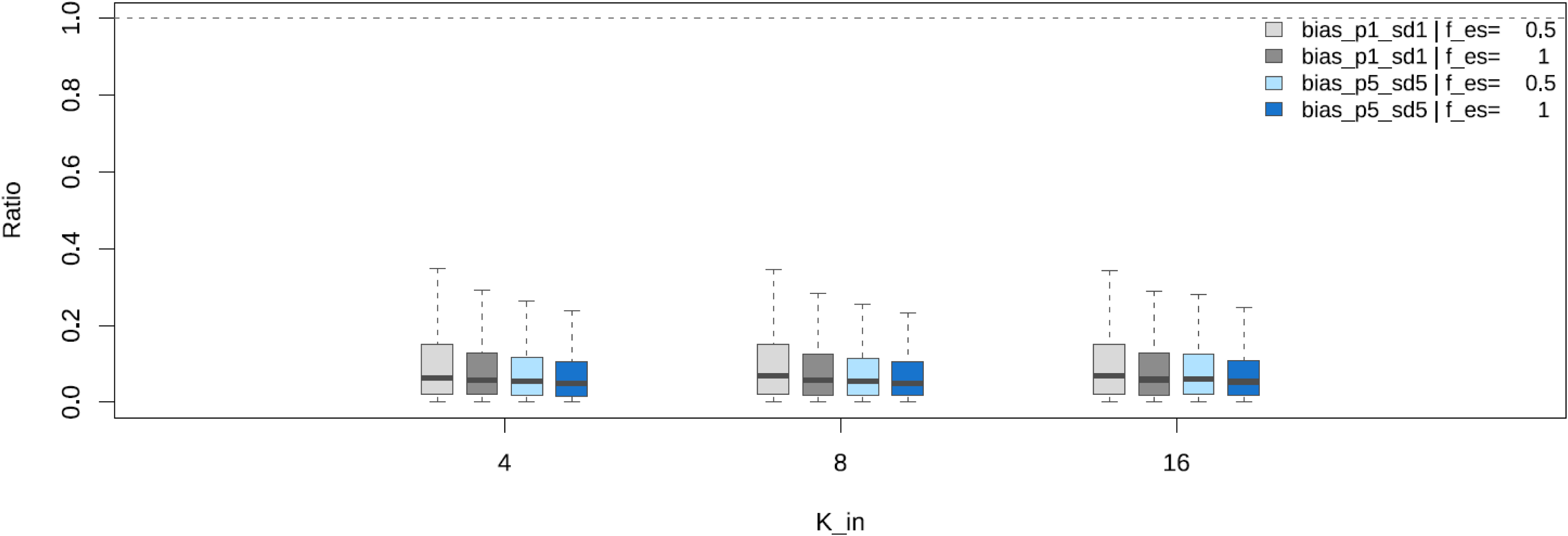
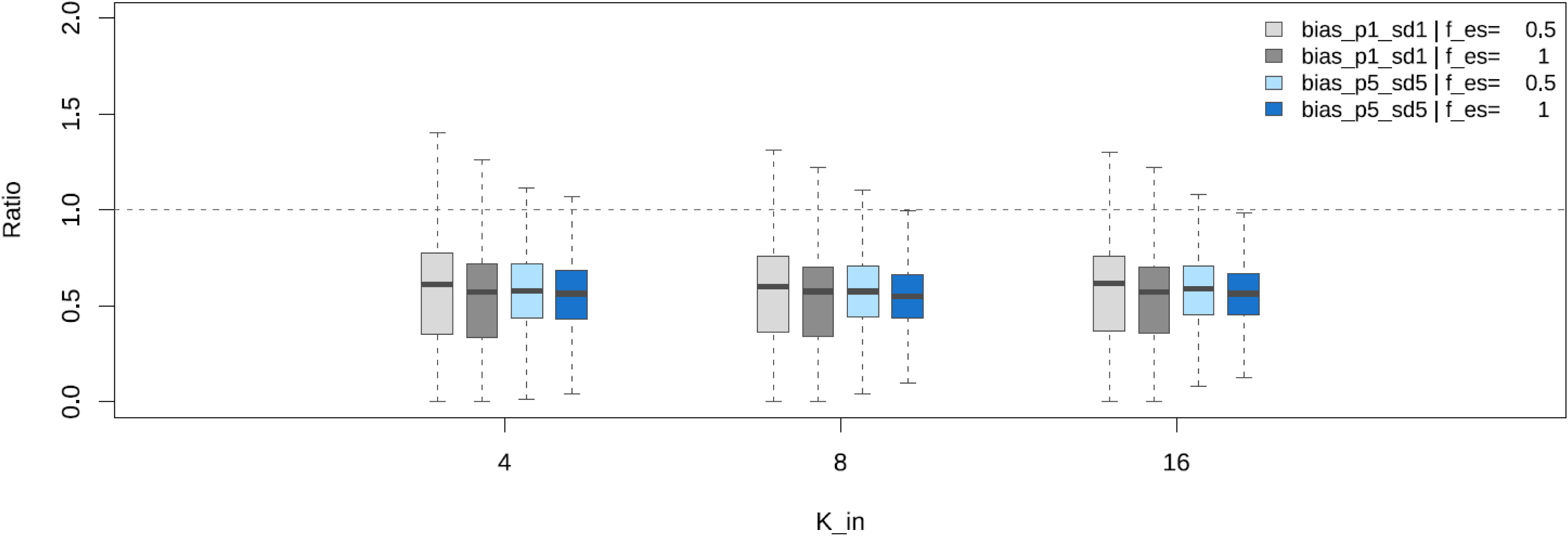
Correction performance in the simulation analysis. (a) Relative bias (RB) after correction, (b) RB ratio calculated using absolute relative bias values, and (c) root mean square error (RMSE) ratio. Results are grouped by error scenario (small– and large-error scenarios), proportion of estimation data (*f*_es_ = 0.5 or 1), and number of cross-validation folds (*K*). Boxplots represent the distributions across bootstrap replicates. Boxes indicate the interquartile range, horizontal lines indicate medians, and whiskers extend to the most extreme values within 1.5 times the interquartile range.

The RB ratio, calculated using absolute RB values, was below one in its central tendency across all conditions. The median was approximately 0.063 in the small-error scenario and 0.053 in the large-error scenario, corresponding to a substantial reduction in the magnitude of RB. In particular, under the large error scenario, improvements were observed across all values of *K*, with slightly better performance when *f*_es_ = 1 compared with *f*_es_ = 0.5 (Fig. 5b).

The RMSE, which reflects agreement in distributional shape, also decreased after correction. The median RMSE ratio was approximately 0.59 and 0.57 in the small-error and the large-error scenario, respectively, corresponding to reductions of approximately 41% and 43%, respectively. In all conditions, the central tendency of the RMSE ratio was below one, indicating that standardization improved the overall shape of the length-frequency distributions (Fig. 5c).

Linear mixed models further clarified the relative effects of the simulation settings (Table 2). For the RMSE ratio, the estimation-data category *F* showed the largest effect among the fixed effects, whereas the effect of the cross-validation-setting category *W* was comparatively small. For the RB ratio, both the error-scenario category *E* and the estimation-data category *F* showed clear effects, whereas *W* showed no clear effect. These results indicate that the amount of paired data used to estimate *P* had a greater influence on correction performance than the number of cross-validation folds. Overall, the RB converged to near zero after correction, and compared with the uncorrected case, both the magnitude of RB and RMSE were reduced in their central tendency. These trends were maintained even under the large error scenario, indicating that, within the range considered in this study, the proportion of estimation data had a greater influence on standardization performance than the number of cross-validation folds.

**Table 2.**
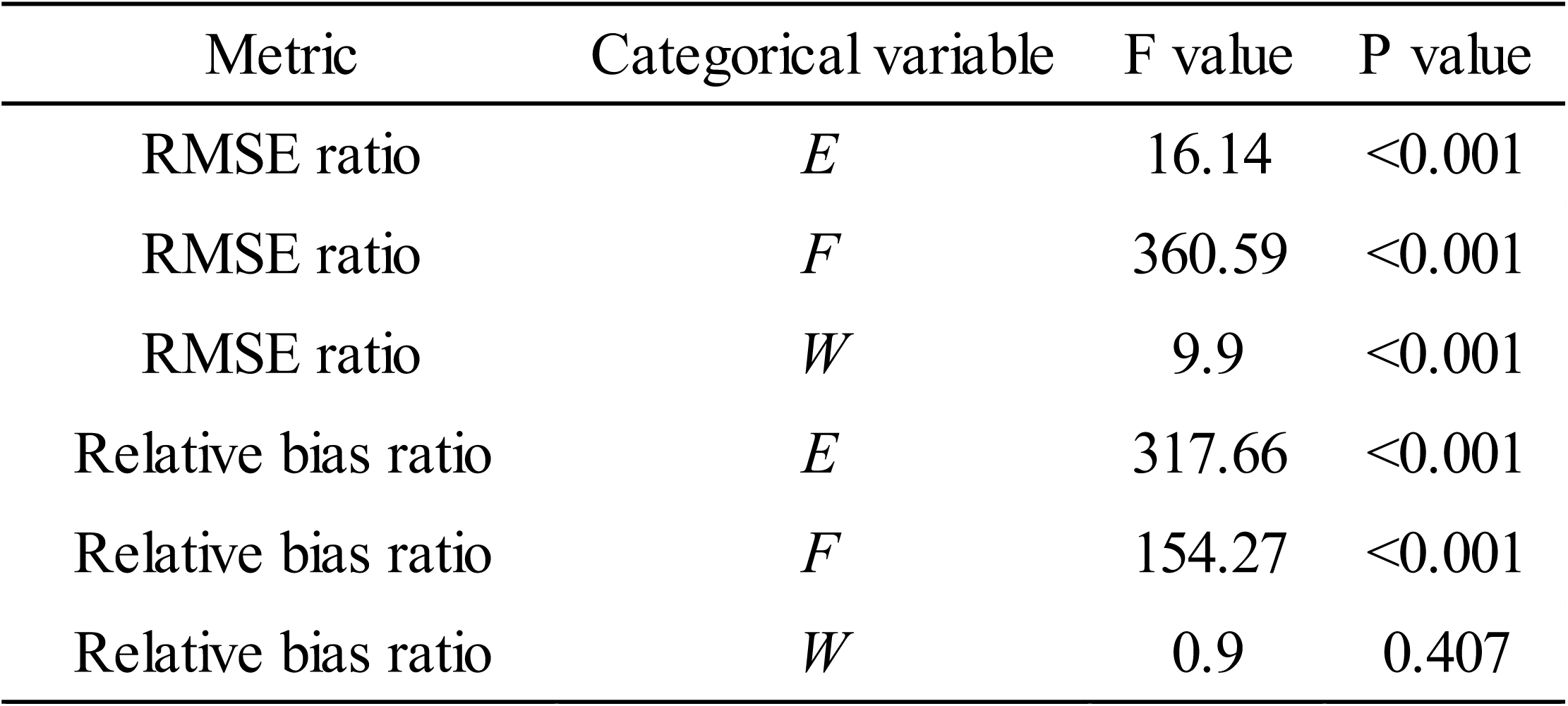
Effects of simulation settings on correction performance based on linear mixed models. The relative bias ratio was calculated using absolute relative bias values.

| Metric | Categorical variable | F value | P value |
| --- | --- | --- | --- |
| RMSE ratio | *E* | 16.14 | <0.001 |
| RMSE ratio | *F* | 360.59 | <0.001 |
| RMSE ratio | *W* | 9.9 | <0.001 |
| Relative bias ratio | *E* | 317.66 | <0.001 |
| Relative bias ratio | *F* | 154.27 | <0.001 |
| Relative bias ratio | *W* | 0.9 | 0.407 |

As an illustrative example, for one species (*Scomber scombrus*) from the ICES DATRAS dataset under the large-error scenario with *f_es_* = 0.5 and *K* = 8, the optimal *m*\* selected via cross-validation was 38 (Fig. 6a). When comparing the histograms before and after correction, the AI-derived distribution clearly approached the reference distribution after standardization (Fig. 6b). In this case, RB decreased from 24.5% to 0.63%, corresponding to an RB ratio of 0.026. The RMSE of the composition probability vectors, evaluated on the evaluation dataset, decreased from 0.02137 to 0.00812, corresponding to an RMSE ratio of 0.38. These results demonstrate that the proposed method improved both the histogram shape and the mean length in this representative case.

**Fig. 6.**
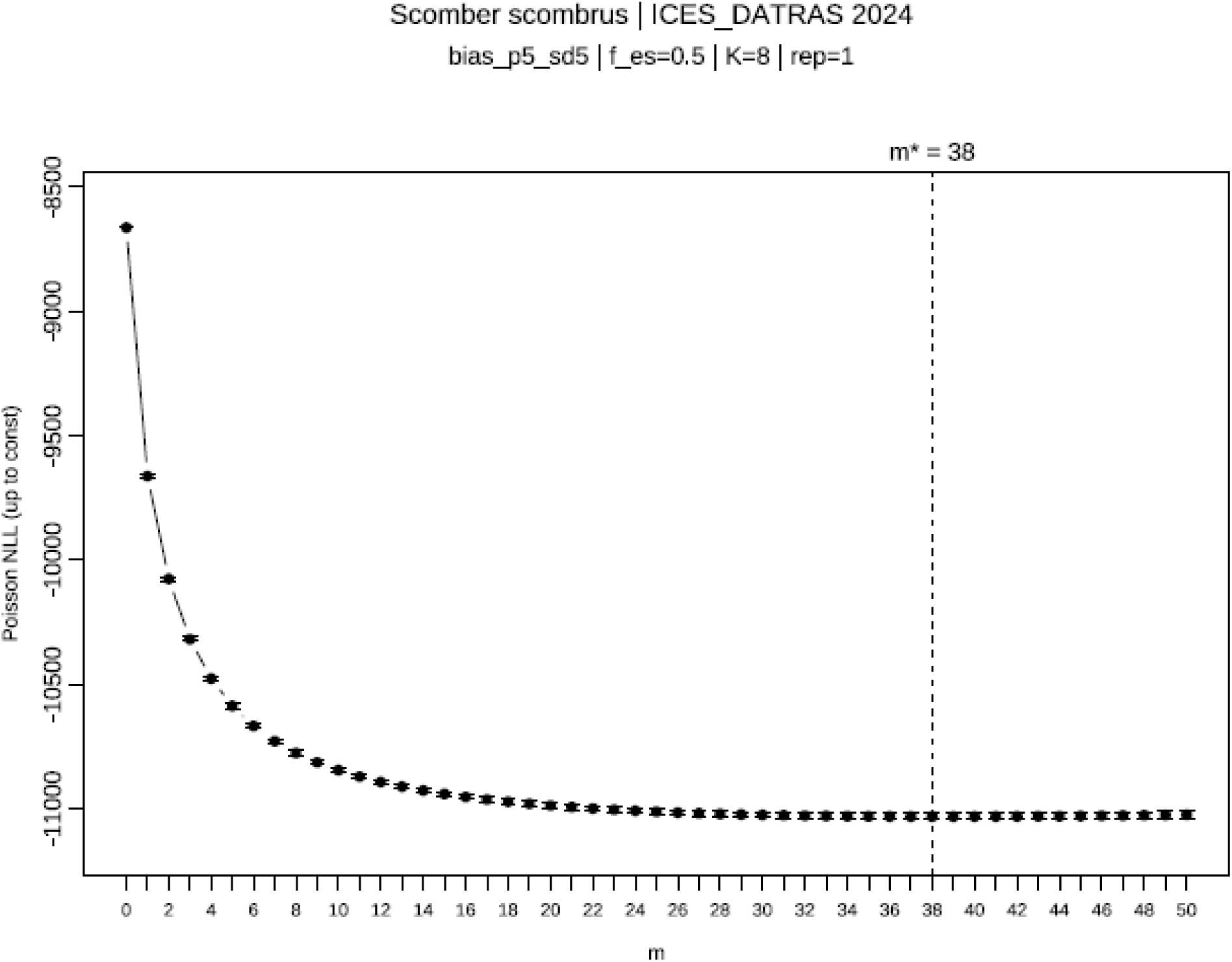
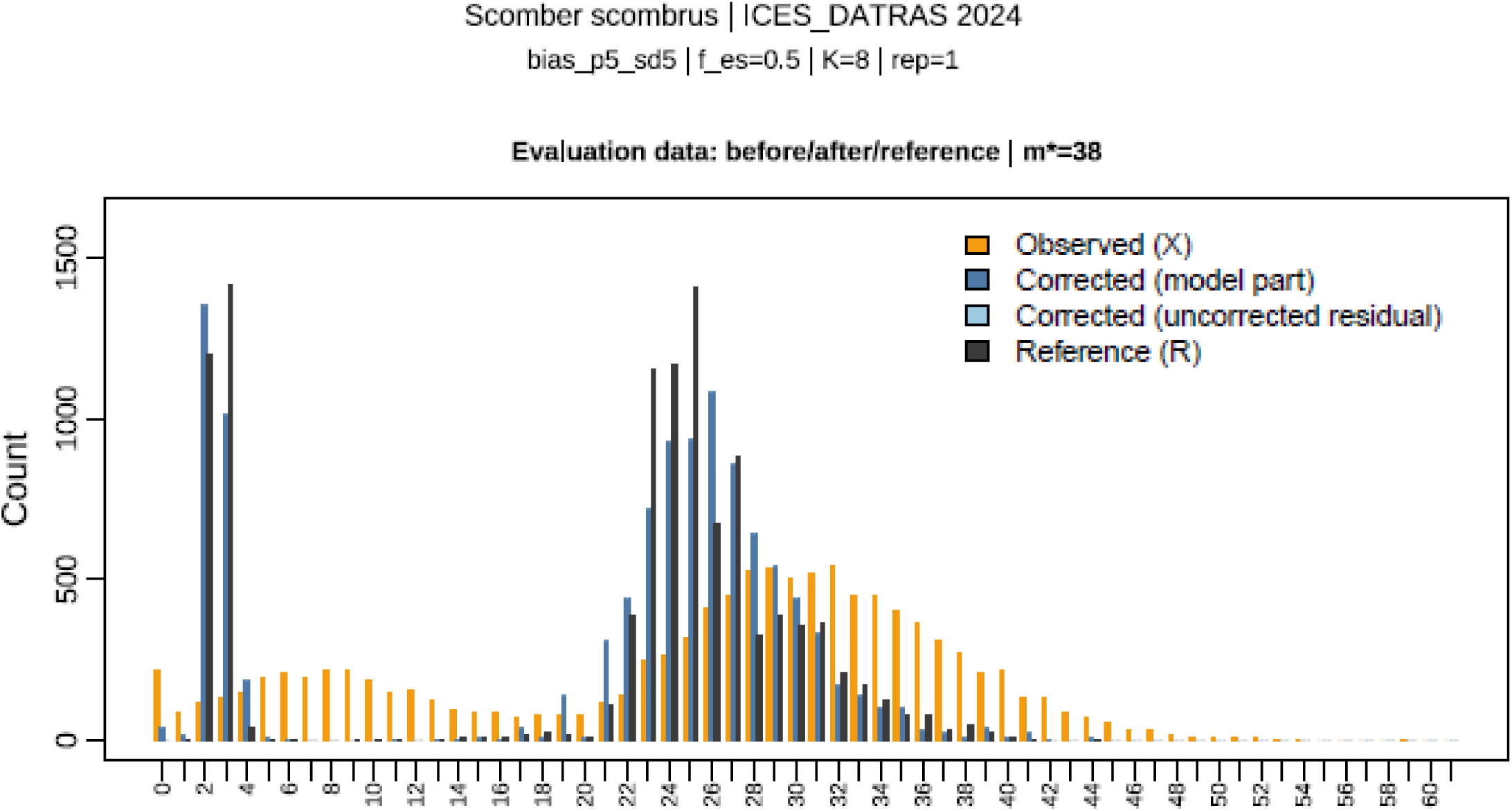
Representative simulation result for one species (*Scomber scombrus*) from the ICES DATRAS dataset. (a) Selection of the optimal number of iterations *m*\* via cross-validation (y-axis: Poisson negative log-likelihood; x-axis: *m*). (b) Comparison of histograms before and after standardization.

### Real-data analysis

In the real-data analysis, the RB of the uncorrected AI-derived histogram showed systematic deviations across length ranges relative to the mean FL of the reference histogram *L*^ˉ^ (*r*_ABC_). The three human observers also showed measurable differences in RB and RMSE, indicating that the reference measurements themselves included inter-observer variability (Fig. 7; Table 3). Following correction using *P*_A_, *P*_B_, and *P*_C_, the AI-derived length-frequency distributions moved closer to the range defined by the three human observers (A, B, and C) and the relative differences in mean FL were substantially reduced (Fig. 7a).

**Fig. 7.**
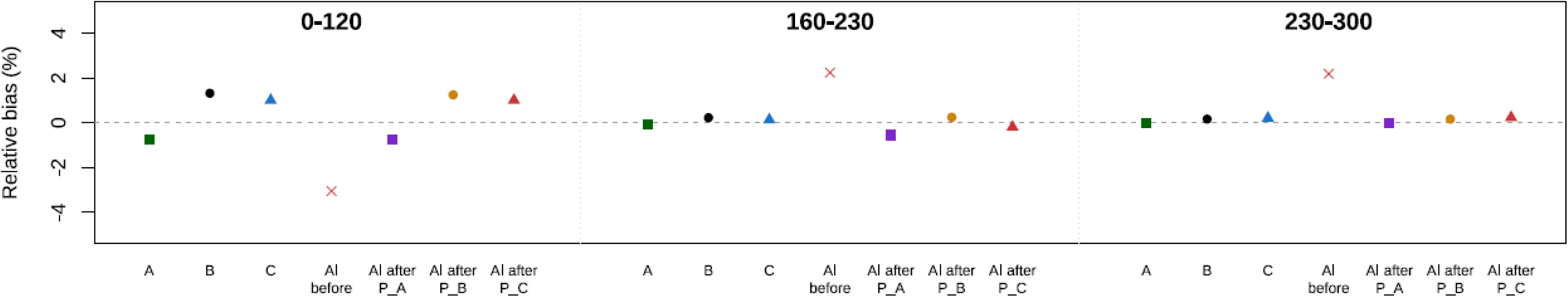
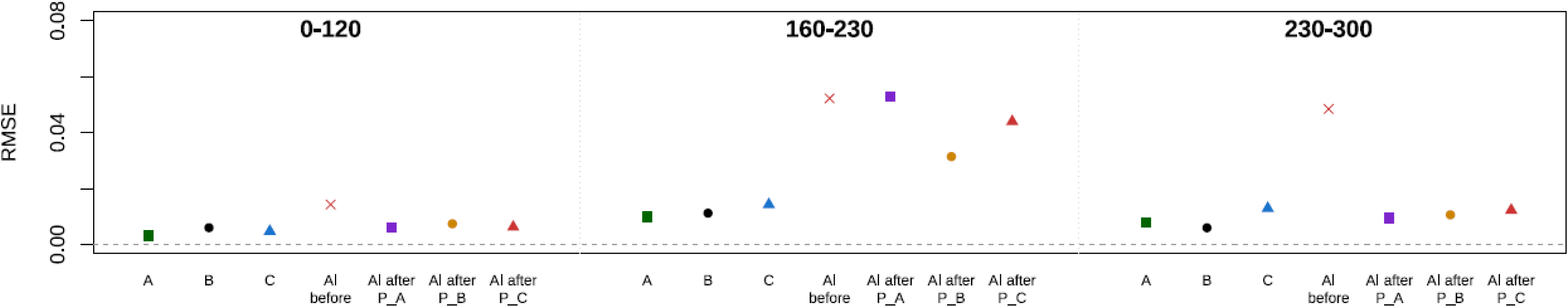
Degree of standardization in the real-data analysis. (a) Relative bias in mean fork length (FL) for each method. (b) Root mean square error (RMSE). The labels 0–120, 160–230 and 230–300 indicate FL ranges in mm.

**Table 3.** Bias and root mean square error (RMSE) of human-observer and AI-derived histograms, including before– and after-correction results using different error matrices (*P*_A_, *P*_B_, and *P*_C_), evaluated across multiple length ranges.

| Observer/AI | Bias |  |  |  | RMSE |  |  |  | RMSE Ratio |  |  |  |
| --- | --- | --- | --- | --- | --- | --- | --- | --- | --- | --- | --- | --- |
|  | All range | 0-120 | 160-230 | 230-300 | All range | 0-120 | 160-230 | 230-300 | All range | 0-120 | 160-230 | 230-300 |
| A | -0.281 | -0.733 | -0.053 | -0.023 | 0.002 | 0.003 | 0.010 | 0.008 |  |  |  |  |
| B | 0.586 | 1.317 | 0.227 | 0.161 | 0.003 | 0.006 | 0.011 | 0.006 |  |  |  |  |
| C | 0.466 | 1.008 | 0.134 | 0.207 | 0.002 | 0.005 | 0.014 | 0.013 |  |  |  |  |
| AI_before | 0.366 | -3.058 | 2.243 | 2.190 | 0.008 | 0.014 | 0.052 | 0.048 | 1.000 | 1.000 | 1.000 | 1.000 |
| AI_after_ $P_A$ | -0.424 | -0.739 | -0.545 | -0.001 | 0.006 | 0.006 | 0.053 | 0.009 | 0.664 | 0.436 | 1.012 | 0.196 |
| AI_after_ $P_B$ | 0.567 | 1.247 | 0.245 | 0.162 | 0.004 | 0.007 | 0.031 | 0.011 | 0.500 | 0.517 | 0.601 | 0.219 |
| AI_after_ $P_C$ | 0.379 | 1.007 | -0.192 | 0.240 | 0.005 | 0.006 | 0.044 | 0.012 | 0.582 | 0.436 | 0.841 | 0.253 |

The RMSE was also higher for the uncorrected AI-derived histogram than for the human-observer histograms. Following the correction, the RMSE decreased markedly. Among the three standardized histograms, *P*_B_ showed the closest agreement with the common reference histogram *r*_ABC_ in terms of overall RMSE, primarily due to improved agreement in the 160–230 mm range (Fig. 7b). The overlaid histograms in Fig. 8 show how the AI-derived histogram changed when each human observer was used as the reference for estimating *P*. They also provide a visual comparison of the resulting standardized histograms with the three human-observer histograms. In this comparison, the standardized histograms were generally closer to the human-observer histograms than the uncorrected AI-derived histogram, although the degree of agreement differed among *P*_A_, *P*_B_ and *P*_C_.

**Fig. 8.**
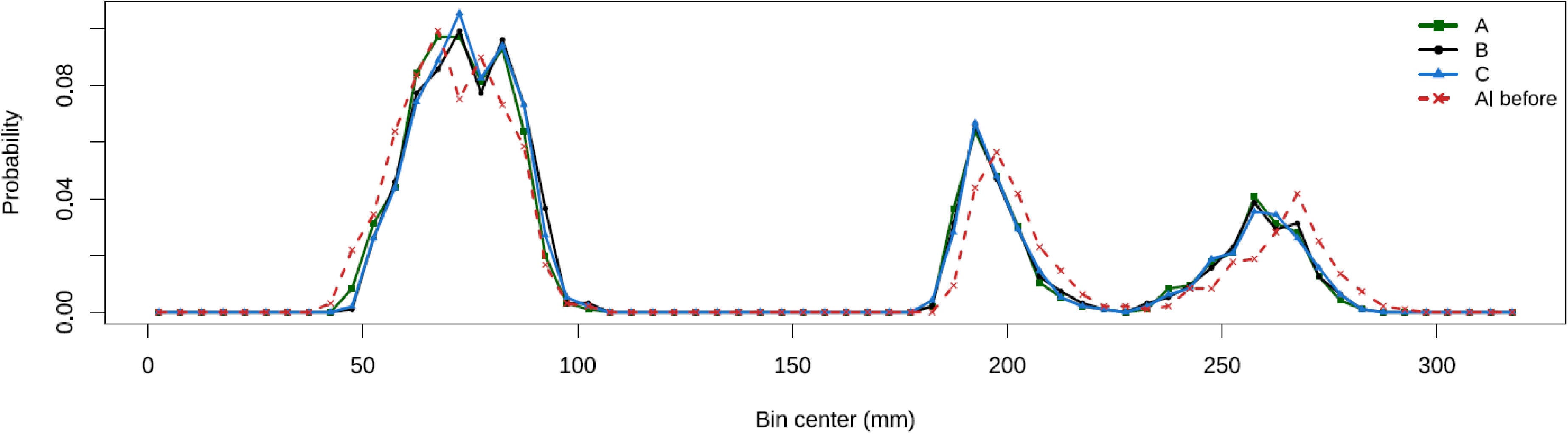
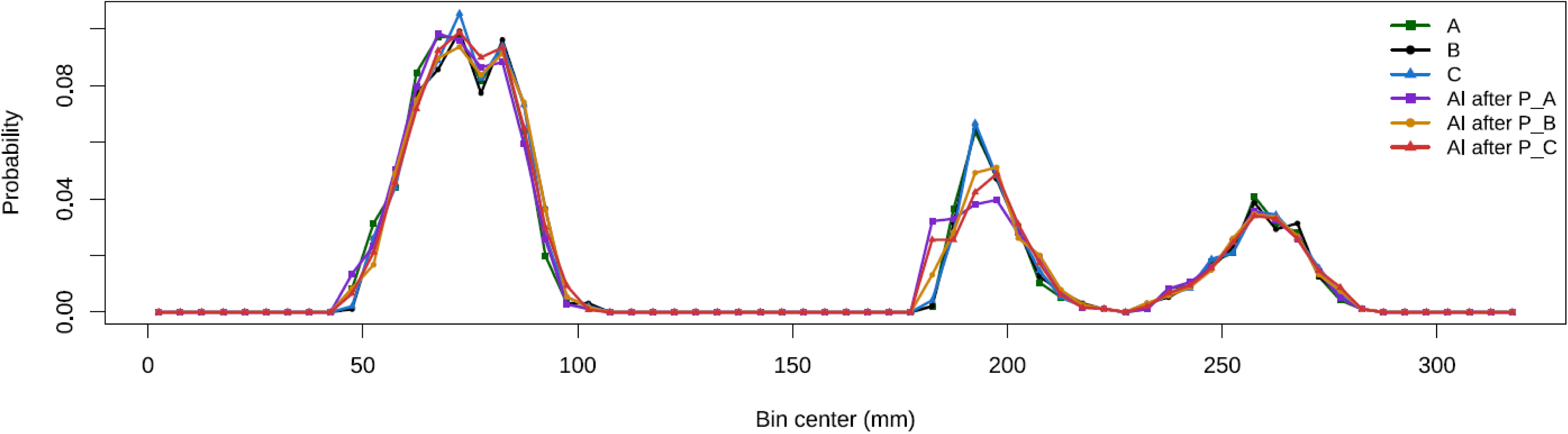
Comparison of length-frequency compositions before and after standardization in the real-data analysis. (a) Comparison between human-observer histograms (A, B, and C) and the uncorrected AI-derived histogram. (b) Comparison between human-observer histograms and AI-derived histograms standardized using observer-specific error matrices (*P*_A_, *P*_B_, and *P*_C_). Human-observer histograms show the candidate reference standards and the range of inter-observer variability. The standardized AI-derived histograms show the results obtained when each observer-specific error matrix was used for standardization.

The superiority of *P*_B_ was also evident from the behavior of bin-wise residuals in the 160–230 mm range (Fig. 9). Although *P*_A_, *P*_B_, and *P*_C_ exhibited broadly similar patterns of positive and negative deviations relative to the reference histogram *r*_ABC_, *P*_B_ suppressed the large oscillations observed in the range of 182.5–192.5 mm more effectively than *P*_A_ and *P*_C_ (Fig. 9b). In other words, the advantage of *P*_B_ did not arise from a fundamentally different correction direction, but from reducing the amplitude of residual fluctuations near local peaks. This resulted in a lower RMSE in the 160–230 mm range, and, consequently, the best overall RMSE among the three candidates (Table 3).

**Fig. 9.**
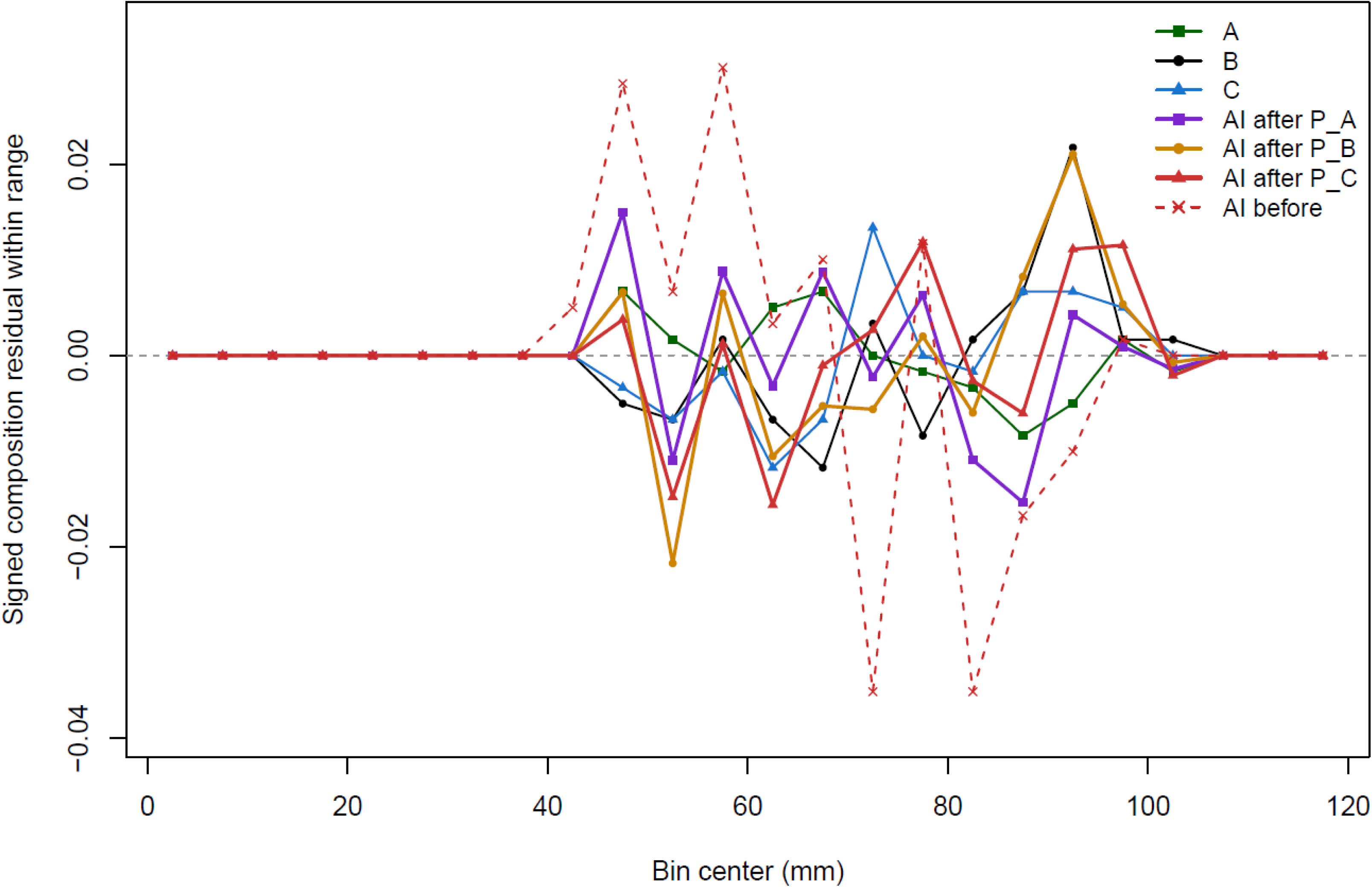
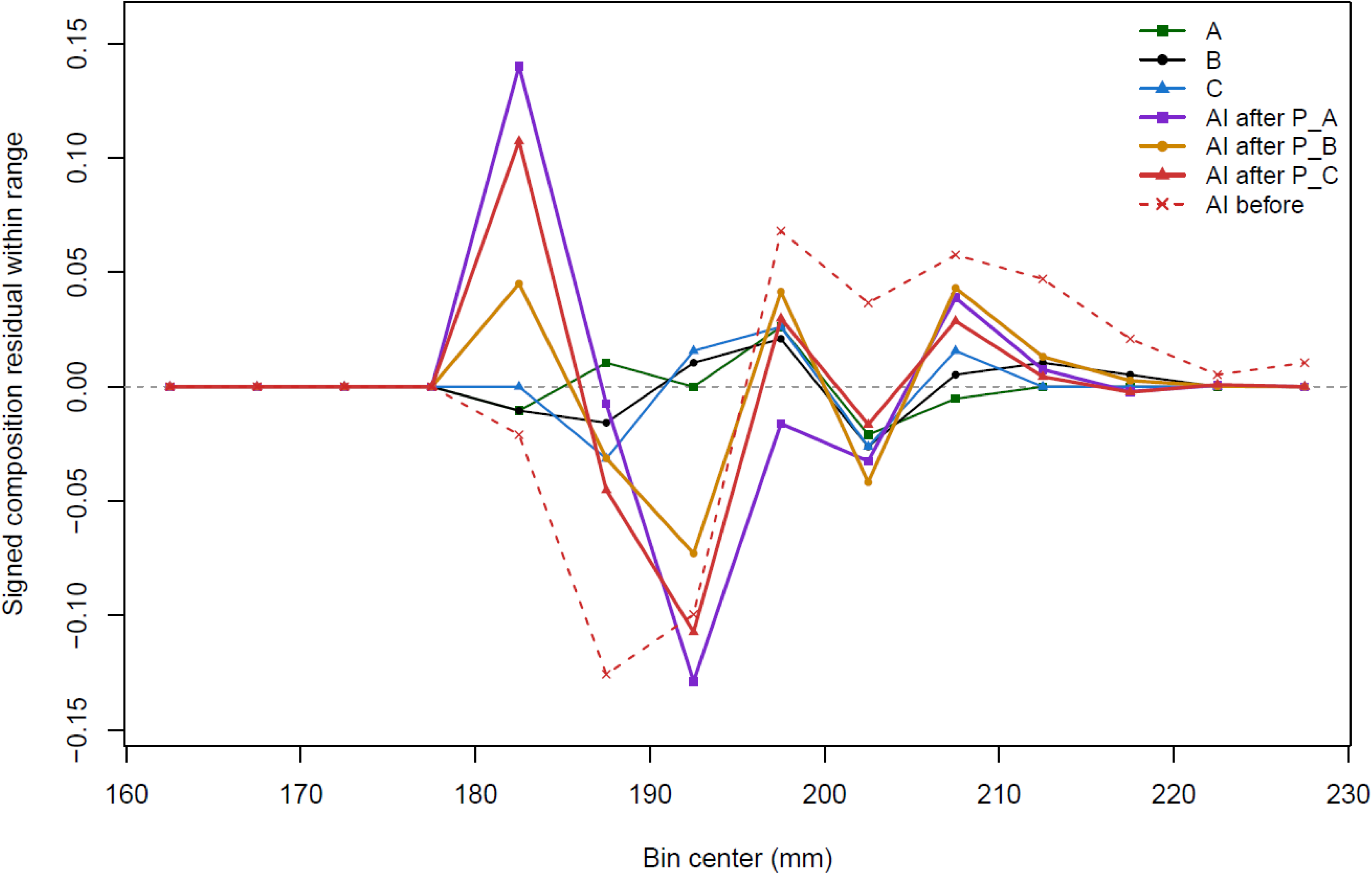
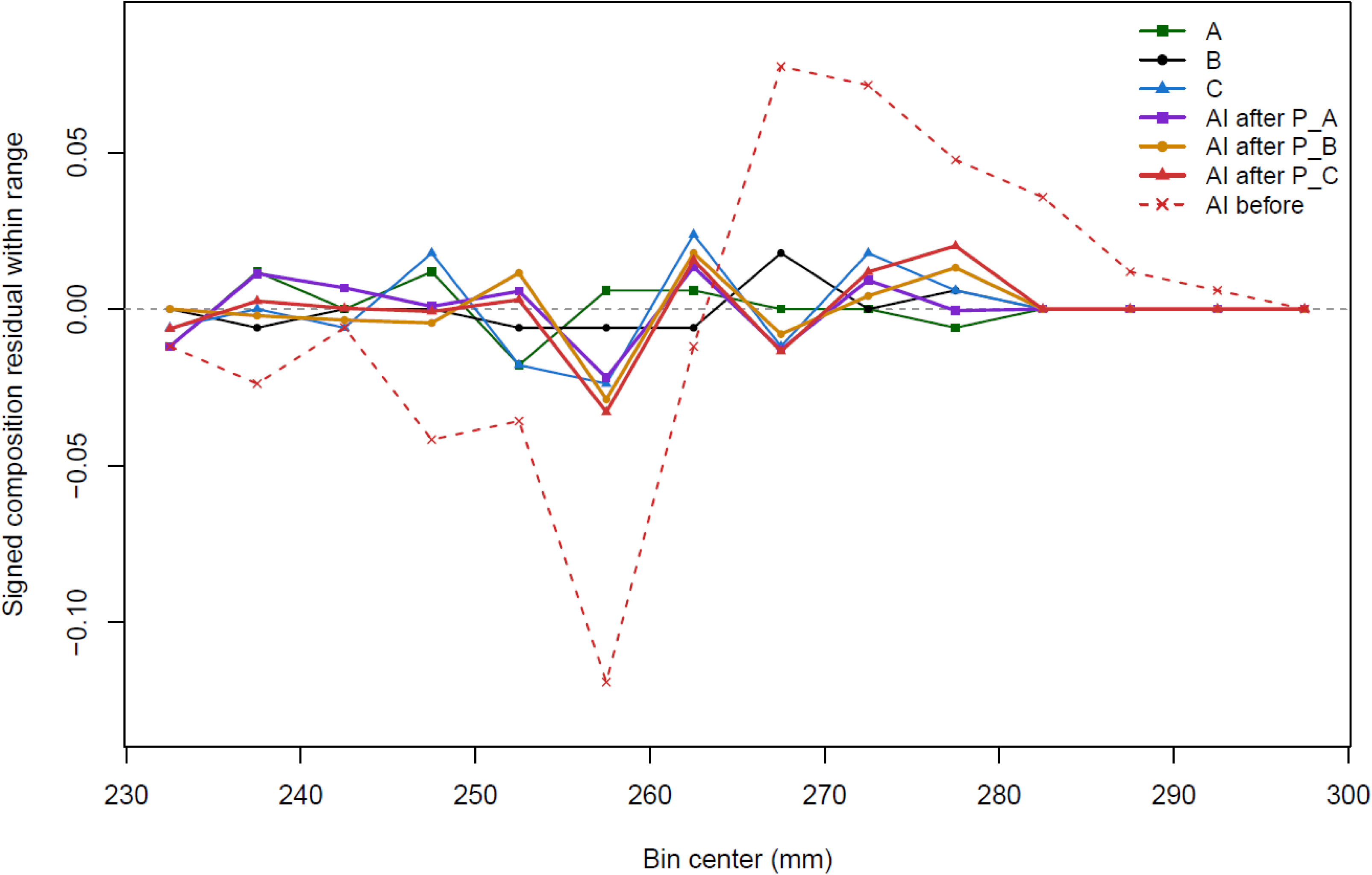
Signed composition residuals between each method and the reference histogram (r_ABC_) in the real-data analysis. For each panel, histograms were restricted to the corresponding length range and converted to composition probability vectors within that range before residuals were calculated. Residuals were calculated as the difference between the composition probability of each method and the reference composition. Residuals for the (a) 0–120 mm range, (b) 160–230 mm range, and (c) 230–300 mm range. Positive values indicate overestimation relative to the reference histogram, whereas negative values indicate underestimation.

## Discussion

Our study demonstrates that a bin-specific error matrix *P* estimated from paired observations can substantially reduce discrepancies between AI-derived and reference-based length-frequency distributions under the examined conditions. A key novelty of the proposed framework is that the estimation of the error structure is separate from the application of the correction. Once a survey-specific *P* is estimated from paired observations, it can be applied as a post hoc correction without retraining the underlying AI model, provided that paired calibration data are available for the target survey. This differs from approaches that seek to improve individual-level prediction by retraining AI models, which often require additional annotation, validation, and model development for each new survey condition. The practical differences between the two approaches are summarized in Table 4.

**Table 4.** Comparison of model-retraining and post hoc error-matrix approaches for handling errors in AI-derived lengths.

| Component | Model retraining approach | Proposed framework |
| --- | --- | --- |
| Image acquisition | Required | Required |
| Pairing with reference measurements | Required | Required |
| Annotation for training (e.g., keypoints, bounding boxes) | Required | Required |
| Annotation for retraining (to align model performance across surveys) | Required | Generally not required |
| Error handling | Embedded in model | Explicitly modeled via error matrix $P$ (post-hoc correction) |
| Transferability across surveys | Limited (requires retraining to align model performance across surveys) | High (cross-survey harmonization without model retraining, given paired data) |
| Operational cost | Higher | Lower (no additional annotation for retraining) |
| Scalability | Limited | High (post-hoc harmonization using paired calibration data; model retraining is generally not required) |

However, the effectiveness of this framework depends on the stability and reliability of the estimated *P*. If the paired data used to estimate *P* are insufficient, biased, or not representative of the underlying observation conditions, the resulting correction may be inaccurate or unstable. In particular, situations with sparse observations in certain length bins, strong domain shifts between training and application conditions, or highly variable measurement errors may result in unreliable estimates of *P* and consequently reduced correction performance. Therefore, careful design of the paired sampling scheme and assessment of the robustness of *P* are crucial for practical application. In practice, these risks can be mitigated by ensuring sufficient sample size and coverage across length bins when constructing paired datasets.

The simulation experiments showed that, under a stable error structure, the proposed method reduced both bias and distributional discrepancy across a wide range of empirical length-frequency distributions. Performance was more strongly affected by the amount of paired data used to estimate *P* than by the number of cross-validation folds. This indicates that securing sufficient paired observations across length bins is more important than increasing *K* within the range examined here. However, the simulation employed a simplified error structure that implicitly included length dependence and did not encompass more complex sources of variation, such as fish posture, imaging angle, or species-specific characteristics.

These results also support the validity of selecting the number of RL-EM iterations *m* via cross-validation. In the representative example, a relatively large value of *m*\* was selected. This result suggests that, although RL-EM is known to amplify noise with increasing iterations, cross-validation can select an appropriate *m*\* that balances reconstruction accuracy and stability, even when the optimal value is relatively large. In this framework, regularization is achieved not necessarily by early stopping, but by selecting an iteration number that optimally balances data fidelity and stability based on cross-validation.

In the proposed framework, individuals falling into unexplainable bins were retained as unstandardized residuals. This design is practically robust in that it avoids artificially forcing agreement by overcorrection and instead explicitly separates the standardizable and non-standardizable components. However, when the amount of paired data is insufficient, the proportion of unstandardized residuals may increase, reducing overall performance. Therefore, in practical applications, it is necessary to ensure a sufficient amount of paired data such that the residual component remains within an acceptable range. Furthermore, when only the AI model is updated, *P* can be re-estimated by applying the new model to the existing images in the paired dataset, and no additional data collection is generally required. However, when imaging conditions change, the existing images may no longer represent the new conditions and additional paired data should be collected for recalibration.

RL-EM was adopted as a practical and theoretically consistent method for reconstructing reference-based length-frequency distributions under a Poisson observation model. In principle, direct matrix inversion or related linear deconvolution methods could also be applied when the estimated *P* is nonsingular and well-conditioned. However, linear inverse problems involving rank-deficient or ill-conditioned matrices are sensitive to perturbations in the input data (Hansen 1998). As *P* in the present framework is estimated from finite paired data and can be sparse or nearly singular, unconstrained inversion may amplify sampling noise and produce negative or highly variable estimates. RL-EM was therefore adopted because it preserves non-negativity and total counts under a Poisson observation model, while cross-validation-based stopping provides a practical form of regularization (Shepp and Vardi 1982; Stanley 1995; Prato et al. 2012). Nevertheless, a systematic comparison with alternative deconvolution methods remains a topic for future work.

In the real-data analysis, the uncorrected AI-derived histogram exhibited systematic deviations across length ranges when compared with the reference histogram *r*_ABC_. Following correction using *P*_A_, *P*_B_, and *P*_C_, the resulting distributions moved closer to the range defined by the three human observers (Fig. 8b). In particular, the results for RB indicate that the mean FL of the standardized AI distributions approached the range of inter-observer variability relative to *L*^ˉ^ (*r_ABC_*). These results indicate that the proposed method can reduce discrepancies between AI-derived length-frequency distributions and human measurements. Rather than using raw AI outputs directly, the proposed method corrects AI-derived length-frequency distributions to make them more consistent with reference measurements.

Among the three candidate error matrices, *P*_B_ yielded the best overall RMSE (Table 3). However, this does not imply that *P*_B_ was uniformly optimal across all length ranges. Although *P*_B_ yielded the lowest overall RMSE and the lowest RMSE in the 160–230 mm range, *P*_A_ yielded the lowest RMSE in the 0–120 mm and 230–300 mm ranges, and *P*_C_ was close to *P*_A_ in the 0–120 mm range (Table 3). The results in the 160–230 mm range are particularly important. The signed residual plots showed that *P*_B_ more effectively suppressed large oscillations from 182.5–192.5 mm compared with *P*_A_ and *P*_C_ (Fig. 9). As RMSE is based on squared differences in probabilities, large residual peaks in a small number of bins can strongly influence the overall metric. Therefore, the advantage of *P*_B_ can be interpreted as arising from its ability to reduce the amplitude of local residual peaks near dominant modes, rather than from a fundamentally different correction direction.

Among the three observers, observer A showed the smallest deviation in mean FL from *L*^ˉ^ (*r_ABC_*) (Table 3). Nevertheless, the best correction performance was obtained using *P*_B_. This indicates that the observer with the smallest mean FL deviation from the consensus reference does not necessarily provide the optimal standardization reference. Instead, the critical factor is which *P* yields smaller discrepancies from the reference histogram in terms of RMSE and residual patterns. In this dataset, *P*_B_ provided the most stable result from this perspective. However, this conclusion is based on consistency within the same dataset and does not imply general superiority for independent datasets.

Meanwhile, even with *P*_B_, RMSE in the 160–230 mm range did not decrease to the level observed among human observers (Fig. 7b). This suggests that, although the mean FL can be corrected effectively, it is more difficult to fully reproduce detailed distributional features, such as the height and width of narrow local peaks using a single error matrix. This difficulty may also be related to the fact that this length range showed lower agreement even among human observers (Fig. 7b). Future work should consider increasing the amount of paired data in this range and exploring standardization approaches that place greater emphasis on local shape reconstruction.

## Practical implications

In practical applications, paired calibration data should cover the dominant modes and tails of the target length-frequency distribution. If multiple reference measurers are available, candidate error matrices can be compared using RB, RMSE, and residual patterns. When independent validation data are available, the matrix that most consistently reduces these discrepancies should be selected. Recalibration is recommended when imaging conditions change or when the existing paired data no longer represent the new operational conditions. Although this study focused on length-frequency data, the same framework may be extended to other classification problems where a confusion matrix can be estimated from paired reference and predicted categories (Spence et al. 2025).

## Data Availability Statement

The processed length-frequency datasets, AI-derived lengths, reference measurements, simulation outputs, and R scripts used to reproduce the analyses and figures in this study have been deposited in an embargoed Zenodo repository (DOI: 10.5281/zenodo.20587024). The repository will be made publicly available upon acceptance of the manuscript. Reviewer access can be provided during peer review upon request.

The processed length-frequency datasets were derived from multiple publicly available data sources and were aggregated and processed for the purposes of this study. They are not intended to serve as substitutes for the original datasets. The original data remain subject to the data policies of their respective providers. ICES DATRAS and RLS data are distributed under the Creative Commons Attribution 4.0 (CC BY 4.0) license, whereas NOAA MRIP and WCPFC data are subject to the respective data usage policies of each organization.

The image dataset used to train the keypoint estimation model and raw images used for the real-data AI inference are not publicly available. However, the AI-derived fork lengths used as fixed inputs in the present analysis, the corresponding reference measurements, and all code required for the downstream standardization analyses have been deposited in the embargoed Zenodo repository.

## Acknowledgment

The authors are grateful to Akiko Niitsuma, Mayumi Suzuki, Hirohito Miyashita, Mikio Watai, Masahiro Manano for their assistance with fish measurements, and to Tohya Yasuda for providing the facilities and experimental environment used in the real-data analysis. The authors also used ChatGPT (OpenAI, GPT-5.5 Thinking) as a tool to assist with code formatting, debugging in R and English language editing. All outputs were critically reviewed and verified by the authors, who take full responsibility for the analyses, interpretations and final manuscript. This study was funded by the Fisheries Agency of the Ministry of Agriculture, Forestry and Fisheries of Japan.

