## Supplementary1 for "Standardizing image-derived fish length-frequency distributions to reference measurements using bin-specific error matrices"

**Supplementary 1. Numerical example: Standardization using an error matrix when unexplainable bins are present**

This supplement provides a small numerical example showing how an error matrix (confusion matrix) is constructed from estimation data, and how Richardson–Lucy expectation–maximization (RL-EM) standardization is performed when the evaluation data include an unexplainable bin. The aim of this example is not to provide a compact mathematical derivation, but to help readers understand how the standardized component and the unstandardized component are separated.

Following the notation in the main text, let $R$ denote the reference length bin and $X$ denote the AI-predicted length bin. The error matrix $P$ is defined as follows, as in Eq. (5) of the main text.

$$P_{i,j} = \Pr(X = i \mid R = j), \qquad \text{(S1)}$$

where columns correspond to reference bins $R$, and rows correspond to AI-predicted bins $X$.

**1. Constructing the error matrix from estimation data**

Suppose that the following paired data are obtained in the estimation dataset.

**Table S1.1. Example of paired reference measurements and AI predictions in the estimation data**

| Fish ID | $R$(cm) | $X$(cm) |
|---|---|---|
| 1 | 20 | 20 |
| 2 | 20 | 20 |
| 3 | 20 | 21 |
| 4 | 21 | 20 |
| 5 | 21 | 21 |
| 6 | 21 | 21 |

In this example, the reference measurements included in the estimation data are only 20 cm and 21 cm. No fish with a reference length of 22 cm is included. Therefore, the reference bins to be standardized are 20 cm and 21 cm. The paired data can be summarized as the following frequency table.

**Table S1.2. Frequency table constructed from the estimation data**

| $X\backslash R$ | 20 | 21 |
|---|---|---|
| 20 | 2 | 1 |
| 21 | 1 | 2 |
| 22 | 0 | 0 |

For simplicity, only the columns corresponding to the supported reference bins, 20 cm and 21 cm, are shown. The row for the AI-predicted 22 cm bin is retained because this bin can appear in the AI-derived histogram. Normalizing each column by its column sum, as in Eq. (6) of the main text, gives the error matrix

$$P = \begin{pmatrix} 2/3 & 1/3 \\ 1/3 & 2/3 \\ 0 & 0 \end{pmatrix}. \qquad \text{(S2)}$$

This matrix means, for example, that a fish with a reference length of 20 cm is predicted by the AI as 20 cm with probability $2/3$, as 21 cm with probability $1/3$, and as 22 cm with probability 0. Similarly, a fish with a reference length of 21 cm is never predicted as 22 cm in this estimation dataset. Thus, according to this error matrix, an AI-predicted value of 22 cm cannot arise from the supported reference bins 20 cm and 21 cm.

**2. Appearance of an unexplainable bin in the evaluation data**

Next, suppose that in another evaluation dataset, the AI-derived histogram is

$$O = \begin{pmatrix} 3 \\ 2 \\ 4 \end{pmatrix}, \qquad \text{(S3)}$$

where the three rows correspond to AI-predicted bins 20 cm, 21 cm, and 22 cm, respectively. This means that the AI predicted 3 fish as 20 cm, 2 fish as 21 cm, and 4 fish as 22 cm. For clarity in this supplement, *O* denotes the full AI-derived histogram before separating unexplainable bins, whereas *o* denotes the explainable component used as input to RL-EM. Here, the estimation data contained reference bins only for 20 cm and 21 cm. Therefore, the target reference-bin set is

$$S = \{20,21\}. \qquad \text{(S4)}$$

Let *S* denote the set of supported reference bins included in the estimation data. According to the definition of unexplainable bins in the main text, an AI bin *i* is unexplainable if

$$\sum_{j \in S} P_{i,j} = 0. \qquad \text{(S5)}$$

For the AI-predicted bin 22 cm, the corresponding row of *P* in Eq. (S2) is

$$(0,\ 0).$$

Therefore,

$$\sum_{j \in S} P_{i,j} = 0, \qquad \text{(S6)}$$

holds for the AI-predicted 22 cm bin. In other words, under the error matrix estimated from the estimation data, the supported reference bins 20 cm and 21 cm cannot generate any AI-predicted fish in the 22 cm bin.

However, the evaluation data contain 4 fish in the AI-predicted 22 cm bin. Therefore, this bin is treated as an unexplainable bin. In this study, such counts are separated into a component used for RL-EM and a component retained as an unstandardized residual:

$$o = \begin{pmatrix} 3 \\ 2 \\ 0 \end{pmatrix}, U = \begin{pmatrix} 0 \\ 0 \\ 4 \end{pmatrix}. \qquad \text{(S7)}$$

Here, $o$ is used as input to RL-EM, whereas $U$ is the vector of unstandardized residual counts retained in the original AI-predicted bin because it cannot be explained by the estimated error matrix.

#### 3. Standardization using RL-EM

RL-EM is applied only to $o$. The purpose is to estimate the reference-based length composition $t$ for the supported reference bins 20 cm and 21 cm. In this example, the target reference bins are only 20 cm and 21 cm. Therefore, $t$ has two elements. The initial value is set as a uniform distribution:

$$t^{(0)} = \begin{pmatrix} 2.5 \\ 2.5 \end{pmatrix}. \qquad \text{(S8)}$$

The total count is 5, which is equal to the total number of fish in $o$.

#### 3.1 Expected AI histogram under the current estimate

According to Eq. (7) in the main text, the expected AI histogram under the current estimate $t^{(0)}$ is

$$\mu(t^{(0)}) = Pt^{(0)}. \qquad \text{(S9)}$$

Using Eqs. (S2) and (S8),

$$\mu(t^{(0)}) = \begin{pmatrix} 2/3 & 1/3 \\ 1/3 & 2/3 \\ 0 & 0 \end{pmatrix} \begin{pmatrix} 2.5 \\ 2.5 \end{pmatrix} = \begin{pmatrix} 2.5 \\ 2.5 \\ 0 \end{pmatrix}. \qquad \text{(S10)}$$

The expected value for the AI-predicted 22 cm bin is again 0. This confirms that the 22 cm bin cannot be explained by the current error matrix.

#### 3.2 E-step: Allocating AI-bin counts to reference bins

Before proceeding to the E-step, it is useful to clarify the relationship between probabilities and expected counts used in RL-EM. The vector *t* represents expected counts in the reference bins, not probabilities. Therefore, the probability that a randomly selected individual belongs to reference bin $j$ is

$$\Pr(R = j) = \frac{t_j}{\sum_{j'} t_{j'}}. \qquad \text{(S11)}$$

Because the error matrix is defined as (S1), the probability that an individual is assigned to AI-predicted bin *i* is obtained from the law of total probability,

$$\Pr(X = i) = \sum_j \Pr(X = i \mid R = j)\Pr(R = j) = \sum_j P_{i,j} \frac{t_j}{\sum_{j'} t_{j'}}. \qquad \text{(S12)}$$

Multiplying both sides by $\sum_{j'} t_{j'}$, the expected number of individuals in AI-predicted bin *i* is

$$\mu_i(t) = \sum_j P_{i,j}\, t_j. \qquad \text{(S13)}$$

Thus, $\mu_i(t)$ is an expected count, whereas

$$\Pr(X = i) = \frac{\mu_i(t)}{\sum_{j'} t_{j'}}, \qquad \text{(S14)}$$

is the corresponding probability. Using Bayes' theorem,

$$\Pr(R = j \mid X = i) = \frac{\Pr(X=i|R=j)\Pr(R=j)}{\Pr(X=i)}. \qquad \text{(S15)}$$

Substituting Eqs. (S11), (S12), and (S14) into Eq. (S15) gives

$$\Pr(R = j \mid X = i) = \frac{p_{ij} t_j}{\mu_i(t)}. \qquad \text{(S16)}$$

This conditional probability is used in the E-step to allocate the observed counts in AI-predicted bin $i$among the possible reference bins.

According to Eq. (9) in the main text, the conditional probability that a fish in AI-predicted bin $i$ originated from reference bin $j$ is

$$\Pr(R = j \mid X = i) = \frac{p_{i,j} t_j^{(m)}}{\sum_{j'} p_{i,j'} t_{j'}^{(m)}}. \qquad \text{(S17)}$$

Using the initial value $t^{(0)}$ =(2.5, 2.5)$^{\mathrm{T}}$, the conditional probabilities for AI-predicted 20 cm are

$$\Pr(R = 20 \mid X = 20) = \frac{(2/3)\times 2.5}{2.5} = \frac{2}{3}, \qquad \text{(S18)}$$

$$\Pr(R = 21 \mid X = 20) = \frac{(1/3)\times 2.5}{2.5} = \frac{1}{3}. \qquad \text{(S19)}$$

Thus, among the 3 fish in the AI-predicted 20 cm bin, the expected allocation is

$$3 \times \begin{pmatrix} 2/3 \\ 1/3 \end{pmatrix} = \begin{pmatrix} 2 \\ 1 \end{pmatrix}. \qquad \text{(S20)}$$

For AI-predicted 21 cm, the conditional probabilities are

$$\Pr(R = 20 \mid X = 21) = \frac{(1/3)\times 2.5}{2.5} = \frac{1}{3}, \qquad \text{(S21)}$$

$$\Pr(R = 21 \mid X = 21) = \frac{(2/3)\times 2.5}{2.5} = \frac{2}{3}. \qquad \text{(S22)}$$

Thus, among the 2 fish in the AI-predicted 21 cm bin, the expected allocation is

$$2 \times \begin{pmatrix} 1/3 \\ 2/3 \end{pmatrix} = \begin{pmatrix} 2/3 \\ 4/3 \end{pmatrix}. \qquad \text{(S23)}$$

Therefore, the expected allocation table for the first E-step is

$$\hat{z}^{(0)} = \begin{pmatrix} 2 & 1 \\ 2/3 & 4/3 \end{pmatrix}, \qquad \text{(S24)}$$

where rows correspond to AI-predicted bins 20 cm and 21 cm, and columns correspond to reference bins 20 cm and 21 cm. This table means that, from the 3 fish in the AI-predicted 20 cm bin, 2 fish are expected to originate from reference 20 cm and 1 fish from reference 21 cm. Similarly, from the 2 fish in the AI-predicted 21 cm bin, 2/3 fish are expected to originate from reference 20 cm and 4/3 fish from reference 21 cm.

#### 3.3 M-step: Updating the reference-based length composition

According to Eq. (11) in the main text, the reference-based length composition is updated by summing the expected counts assigned to each reference bin over all AI bins. Thus, after the first update,

$$t^{(1)} = \begin{pmatrix} 2 + 2/3 \\ 1 + 4/3 \end{pmatrix} = \begin{pmatrix} 8/3 \\ 7/3 \end{pmatrix}. \qquad \text{(S25)}$$

This is the result after one RL-EM iteration. It is not yet the final solution. Repeating the E-step and M-step gradually updates the reference-based composition. In this example, the iteration converges to

$$\hat{t} = \begin{pmatrix} 4 \\ 1 \end{pmatrix}. \qquad \text{(S26)}$$

To check that this solution is consistent with the explainable part of the AI histogram, we multiply it by the error matrix:

$$P\hat{t} = \begin{pmatrix} 2/3 & 1/3 \\ 1/3 & 2/3 \\ 0 & 0 \end{pmatrix} \begin{pmatrix} 4 \\ 1 \end{pmatrix} = \begin{pmatrix} 3 \\ 2 \\ 0 \end{pmatrix}. \qquad \text{(S27)}$$

This reproduces $o$. This point is important. The RL-EM estimate is not $(3, 2)^T$. The vector $(3, 2, 0)^T$ is the AI-space histogram reproduced by applying $P$ to the estimated reference-based composition. The standardized reference-based component is $\hat{t} = (4, 1)^T$.

### 4. Final standardized length composition

In the main text, the final standardized length composition is defined as the sum of the component standardized by RL-EM and the unstandardized component $U$ retained from the unexplainable bins. In this example, the standardized component obtained by RL-EM is $\hat{t} = (4, 1)^T$. Written over the three length bins 20 cm, 21 cm, and 22 cm, this becomes

$$\hat{y}_{model} = \begin{pmatrix} 4 \\ 1 \\ 0 \end{pmatrix}. \qquad \text{(S28)}$$

The unexplainable component is

$$U = \begin{pmatrix} 0 \\ 0 \\ 4 \end{pmatrix}. \qquad \text{(S29)}$$

Therefore, the final standardized length composition is

$$\hat{y}_{total} = \hat{y}_{model} + U = \begin{pmatrix} 4 \\ 1 \\ 0 \end{pmatrix} + \begin{pmatrix} 0 \\ 0 \\ 4 \end{pmatrix} = \begin{pmatrix} 4 \\ 1 \\ 4 \end{pmatrix}. \qquad \text{(S30)}$$

The important point is that the 4 fish in the 22 cm bin are not the result of RL-EM standardization. They remain in the original AI-predicted bin because they could not be explained by the error matrix estimated from the estimation data. Therefore, $\hat{y}_{total}$ consists of two parts: the standardized component estimated by RL-EM and the unstandardized component retained because it was unexplainable.

### 5. Interpretation of this example

In this numerical example, the estimation data contain reference measurements only in the 20 cm and 21 cm bins. As a result, the error matrix constructed from the estimation data cannot generate AI-predicted values in the 22 cm bin. However, the evaluation data contain 4 fish in the AI-predicted 22 cm bin. This bin is therefore treated as an unexplainable bin. In this study, such fish are not forcibly redistributed to other reference bins. Instead, they are retained as the unstandardized component $U$, and RL-EM is applied only to $o$. Thus, in the presence of unexplainable bins, $o$ in the

RL-EM update denotes the explainable component of the AI-derived histogram. This treatment allows the procedure to continue without stopping the computation. At the same time, it makes clear which part of the final length composition was standardized using the error matrix and which part remained unstandardized because it could not be explained by the estimation data.
